# Label-free imaging flow cytometry: analysis and sorting of enzymatically dissociated tissues

**DOI:** 10.1101/2021.05.05.442869

**Authors:** Maik Herbig, Karen Tessmer, Martin Nötzel, Ahsan Ahmad Nawaz, Tiago Santos-Ferreira, Oliver Borsch, Sylvia J. Gasparini, Jochen Guck, Marius Ader

**Affiliations:** Biotechnology Center, Center for Molecular and Cellular Bioengineering, Technische Universität Dresden, Dresden, Germany; Center for Regenerative Therapies (CRTD), Dresden, Germany; Max Planck Institute for the Science of Light & Max-Planck-Zentrum für Physik und Medizin, Erlangen, Germany; Roche Innovation Center Basel, F. Hoffman-La Roche Ltd., Basel, Switzerland

## Abstract

Biomedical research often relies on identification and isolation of specific cell types using molecular biomarkers and sorting methods such as fluorescence or magnetic activated cell sorting. Labelling processes potentially alter the cells’ properties and should be avoided, especially when purifying cells for clinical applications. A promising alternative is the label-free identification of cells based on their physical properties. Sorting real-time deformability and fluorescence cytometry (soRT-FDC) is a microfluidic technique for label-free analysis and sorting of single cells. In soRT-FDC, bright-field images of cells are analyzed by a deep neural net (DNN) to obtain a sorting decision, but sorting was so far only demonstrated for blood cells which show clear morphological differences and are naturally in suspension. Most cells, however, grow in tissues, requiring dissociation before cell sorting which is associated with additional challenges including survival, changes in morphology, or presence of aggregates. Here, we introduce methods for robust analysis and sorting of single cells from mammalian nervous tissue and provide DNNs which are capable of distinguishing visually similar cells. Exemplarily, we employ the DNN for image-based sorting to enrich photoreceptor cells from dissociated retina for transplantation into the mouse eye. Results provide evidence that the combination of machine learning and soRT-FDC allows label-free enrichment of target cells from dissociated tissues.

## Introduction

Cell characterization is a major task in biomedical research as it allows for refined analyses and isolation of specific cell types for characterization or therapeutic applications. The current gold standard for cell-typing relies on the identification of unique proteins expressed by the target cell population. If the cell-specific protein is intracellular, it cannot be accessed in live cells and genetic engineering is required to introduce expression of a fluorescent reporter. If the protein is located on the cell surface, commercially available antibodies allow to label the cells with a fluorescent or magnetic marker. Thus, the target cells become detectable for fluorescent activated cell sorting (FACS) or magnetic activated cell sorting (MACS), respectively ^[1–3]^. However, specific markers for many cell types have still not been defined and such labeling processes present a treatment which could alter the cells’ properties and therefore skew any subsequent analysis or cellular response. Additionally, enzymatic dissociation of tissues frequently affects binding sites essential for the recognition by specific antibodies ^[4]^. A promising alternative is the label-free identification of cells based on inherent physical and morphological properties.

Density gradient centrifugation and filtration-based approaches are well-established methods allowing to enrich cells based on their density and cell size – however these have limited sensitivity. Deterministic lateral displacement is a microfluidic method for the enrichment of cells based on their deformation and size characteristics ^[5]^. Similar to MACS, these techniques sort cells passively, allowing for bulk processing, resulting in an unmatched throughput. On the downside, these bulk sorting techniques only allow to enrich cells based on a small number of characteristics which are often shared by diverse cell types in a tissue and the sorting logic is hard-wired into the setup. In contrast, single cell approaches allow a flexible tuning of the sorting decision. Arguably, the most popular label-free single cell analysis and sorting device is FACS as it allows to obtain size and transparency information (forward scatter and side scatter) without need for staining, however not all cell types can be distinguished using this method ^[6]^. Further inherent properties are chemical composition and mass density, and corresponding methods for single cell analysis or sorting have already been demonstrated ^[7,8]^.

A microfluidic method for capturing mechanical properties of single cells is real-time deformability cytometry (RT-DC), in which cells flow through a narrow channel where they are deformed and captured by a high-speed camera ^[9]^. For retrieval of mechanical features, only the outline of the tracked object (contour) is required, but the technology also provides bright-field images. RT-DC was complemented with fluorescence detection capability (real-time fluorescence and deformability cytometry – RT-FDC ^[10]^), which allows to record bright-field images and fluorescence information of conventional markers simultaneously at a throughput of up to 1,000 cells/s. This technology proved to be ideal to generate labeled image datasets for training deep neural nets (DNNs) which learn to detect cell types based on the bright-field image alone. More recently, a sorting unit was added to the RT-FDC setup (sorting real-time deformability and fluorescence cytometry - soRT-FDC ^[11]^) which leverages real-time image analysis by a DNN to actuate the sorting trigger based on the classification score. The study demonstrated DNN-assisted, image-based sorting of blood cells, which are cells that are naturally occurring in suspension. However, most cells grow in tissues, resulting in a need for dissociation before any kind of single cell flow cytometry method can be applied. The same applies for 2D or 3D cell cultures such as organoids. Organoids are an increasingly popular tool in biomedical research for investigation of developmental and pathologic mechanisms, and they represent a promising cell source for therapeutic purposes ^[12–14]^.

Besides possibilities for label-free sorting, tissue dissociations are subject to non-uniform outcome. While naturally suspended cells tend to show a round shape (e.g. cells from blood or bone marrow), the morphology of cells in tissues is more heterogeneous. Alone in the retina, the shapes range from elongated (e.g. Müller glia) to elliptical (e.g. retinal pigment epithelium), resulting in a broad range of cell morphologies even after dissociation. Furthermore, single cells are prone to aggregate. Aggregates are unfavorable as they could skew a measurement, create artifacts in analysis results, cause accidental sorting of undesired cells, or even congest a sorting unit. Due to the heterogeneous morphologies of dissociated cells and their tendency to aggregate, automatically differentiating single cells and cell aggregates is challenging.

In the present work, we introduce software and hardware methods to improve reliability of RT-FDC data analysis and image-based cell sorting in the context of enzymatically dissociated tissues. We updated the chip design to promote a microfluidic based division of cell aggregates. Furthermore, we trained a convolutional neural net (CNN) for detection of aggregates in images which can be employed for offline analyses of RT-FDC datasets. For real-time detection of aggregates during sorting, we introduce efficient algorithms that employ object counting and the frequency of the occurrence of cells. soRT-FDC was previously demonstrated for DNN-based sorting of blood cells which show prominent phenotype differences ^[11]^. In this work, we describe a DNN architecture for optimized utilization of CPU (central processing unit) resources which improves the accuracy of image-based cell identification for sorting. To demonstrate the applicability of the method to biomedical research, we trained the DNN for detection of photoreceptors from dissociated mouse retinae. The trained model was employed for label-free image-based sorting of photoreceptors, which were subsequently transplanted into adult mice and were successfully shown to survive and interact with the host retina.

## Results

### Hardware based reduction of cell aggregates

In the present work, we build upon the existing soRT-FDC technology to improve reliability of measurement, analysis, and sorting of enzymatically dissociated tissues. To develop and showcase the methods, we used dissociated retina cells originating from human retinal organoids (HROs) and mouse eyes (see Figure 1A). HROs differentiated from a photoreceptor-specific reporter human induced pluripotent stem cell line (hiPSC-Crx-mCherry ^[14]^) were cultured for 125 days. Mice expressing GFP restricted to rod photoreceptors (Nrl-eGFP mouse ^[15]^) were at postnatal day 4 (P04) when applying the dissociation protocol (see Materials and Methods). For flow cytometry measurement in RT-FDC or sorting using soRT-FDC, cells were resuspended in a measurement buffer with elevated viscosity (see Materials and Methods), as illustrated in Figure 1A.

**Figure 1:**
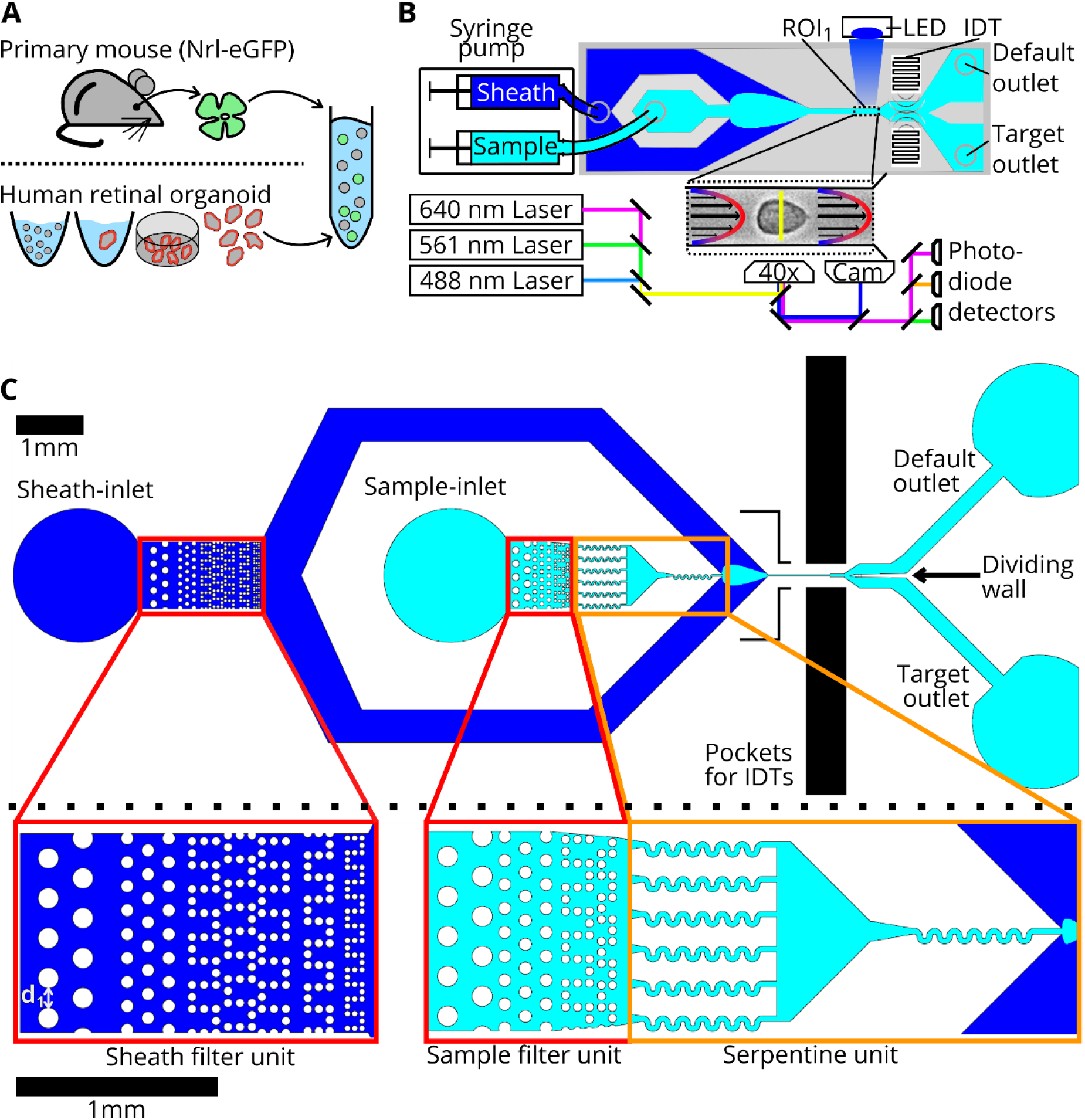
Cell preparation, soRT-FDC setup and chip design (A) Retinae from reporter mice (Nrl-eGFP) or human retinal organoids (Crx-mCherry) are dissociated and resuspended in measurement buffer for soRT-FDC. (B) Sketch of the soRT-FDC setup. Two syringe pumps supply a microfluidic chip with sample and sheath fluid. Lasers excite fluorescence signal which is measured by avalanche photodetectors and the cell is imaged by a high-speed camera. A high-power LED illuminates the cell. Interdigital transducers (IDTs) excite surface acoustic waves, which push selected cells towards the target outlet. (C) Figure shows the 2D-CAD design of the entire sorting chip and zoomed in versions show specific parts. The red rectangles indicate filter assemblies, which consist of a cascade of pillars with decreasing distance. The orange rectangles indicate a unit of several serpentines, which helps to divide aggregates of cells and to increase the spacing between cells. The layout was designed using KLayout 0.25.3.

soRT-FDC is a microfluidic technique allowing not only to capture bright-field images and fluorescence information from single cells at 1,000 cells/s, but also sort specific cells based on the decision of a DNN. In soRT-FDC, suspended cells and sheath fluid are pumped into a microfluidic chip by means of two syringe pumps. The sheath flow focusses the sample flow towards a narrow channel. At the end of the channel the cells are captured by a high-speed camera and optionally fluorescence information is retrieved for up to three wavelengths. After the narrow channel, the microfluidic system widens and divides into a path towards the default and target outlet (see Figure 1B). The narrow channel is a distinct feature of soRT-FDC as it allows to deform cells to obtain information about the mechanical properties of cells. Furthermore, cells are aligned in the channel which simplifies image analysis tasks due to the reduced degrees of freedom.

However, any constriction in a microfluidic design introduces the risk of being blocked by debris or large objects contained in the processed sample. A blocked or partially blocked channel will impair sorting. Moreover, presence of cell clumps in a dataset can skew analysis results. Especially for dissociated samples, presence of cell aggregates like doublets is very common. To prevent such objects from reaching highly confined parts of the chip, multiple columns of filter pillars were implemented. The distance between pillars at the first column is 60 μm (indicated as d1 in Figure 1C), which allows to catch larger objects (see Figure 1C). The pillars at the final column show a distance of 15 μm, which catch smaller objects and also contribute to separating and dividing aggregates into single cells. In the sheath inlet the first and last column of filters have an inner distance of 60 μm and 10 μm, respectively. Separation of cells is further promoted by serpentine channels of a width of 30 μm (see Figure 1C and Figure S1, Supporting Information). We observed that debris particles are prone to get stuck in the curvature of the serpentines. To prevent a full blocking of the chip, multiple serpentines were placed in parallel, resulting in a practically undisturbed execution of measurements or sorting experiments for hours.

The microfluidic design shown in Figure 1C decreases the probability of the occurrence of large aggregates (see Figure S1, Supporting Information) but does not guarantee to generate a pure single cell suspension. In the following, a method for detection of aggregates such as cell doublets is introduced, allowing to exclude such events during data analysis.

### DNN based detection of cell aggregates

In flow cytometry, cell doublets can skew datasets and any subsequent analysis requires an exclusion of such events. For example, when a non-fluorescent cell is attached to a fluorescent cell, the event would be assigned to the fluorescence positive group but other features such as granularity are affected by both cells. Image flow cytometers like RT-FDC and soRT-FDC provide a bright-field image and doublets of cells could be identified by human eye. As datasets typically contain several thousands of images, this task would be extremely labor intensive, resulting in a need for automation. Therefore, we visually assessed more than 60,000 cells (42,583 single cells and 21,137 doublets of cells) using RT-FDC measurements of HROs to create a labelled dataset. To speed up the labelling process, we developed a dedicated software (*YouLabel*) with graphical user-interface (Figure S2, Supporting Information). Using the generated dataset we trained supervised machine learning models, more specifically, convolutional neural nets (CNNs, Figure 2A), a type of DNN that is commonly used for image classification tasks. The input image size for the CNN is 36×36 pixels (= 24.5×24.5 μm) which is large enough to cover aggregates of cells and cells in proximity (Figure 2A). Accidental sorting of multiple cells and erroneous assignment of fluorescence intensities is not only a problem when cells are directly attached to each other but also when they travel at a close distance (see Figure S3 A, Supporting Information). To train the CNN to detect such events, they were assigned to the class of doublets during the manual labeling process.

**Figure 2:**
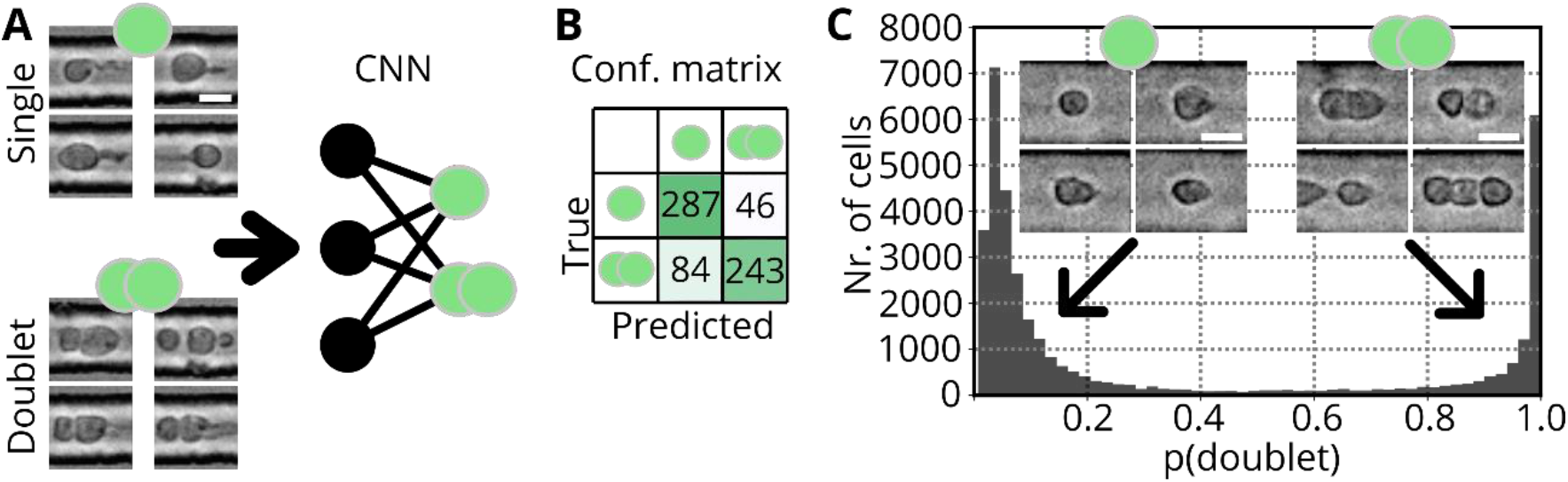
CNN for detection of cell aggregates (A) Bright-field images of single cells and cell aggregates of human retinal organoid cells. Images are used to train a CNN for discrimination between single cells and cell aggregates. (B) Confusion matrix resulting when applying the CNN on the validation set. The validation accuracy is 80.3%. (C) Probability distribution resulting when applying the model to a testing dataset of dissociated Nrl-eGFP retina. Despite the different origin of the cells, the model is able to distinguish between single (left, low probability) and aggregated cells (right, high probability).

In order to span a wide variety of phenotypes, we used images of dissociated HRO cultures ^[16]^. Based on the resulting dataset, we trained a CNN (Figure 2A) to perform the task of identifying doublets, and the resulting model (CNN_doublet_) reaches a validation accuracy of 80.3% (Figure 2B). To test the applicability of the model to new data, we recorded a dataset of murine Nrl-eGFP cells. In Nrl-eGFP transgenic mice GFP expression is restricted to rod photoreceptors ^[15]^. Each event was forwarded through CNN_doublet_ to obtain the probability that the event is a doublet (p_doublet_) and the histogram in Figure 2C shows the resulting distribution of probabilities. Interestingly, the model confidently predicts single cells and doublets into the correct class as shown by example images. The CNN classifies an event as doublet if a second cell is closer than approximately 15 μm (Figure S3 B, Supporting Information). The model also delivers sensible results for a measurement of whole blood (Figure S3 C, Supporting Information, data taken from ^[11]^), indicating that the model could be employed for a general-purpose doublet detection algorithm.

CNN based detection of cell aggregates is a helpful tool for analyzing RT-DC or RT-FDC data which could be employed for many datasets and comes at low computational cost. Forwarding a single image through CNN_doublet_ only requires 1.4 ms (Intel Core i7 3930K @ 3.2 GHz). Processing 10,000 images at once (batch processing) allows to achieve an inference time of 0.75 ms per image. While these times are sufficient to process large datasets, for sorting an inference time below 250 μs is required. Therefore, faster doublet detection methods are required.

### Detection and separation of cell aggregates for single cell sorting

In RT-DC, RT-FDC, and soRT-FDC a real-time contour detection algorithm evaluates acquired images using efficient OpenCV implementations. By counting the number of contours in an image, we implemented a switch that allows to suppress sorting if more than n=1 contours were detected (see Figure 3A). The additional contour counting step comes at no additional computational cost. To reduce the chance of having multiple cells within the ROI, the cell concentration could be decreased but since that would decrease the frequency of measurement and sorting, an optimal cell concentration needs to be determined.

**Figure 3:**
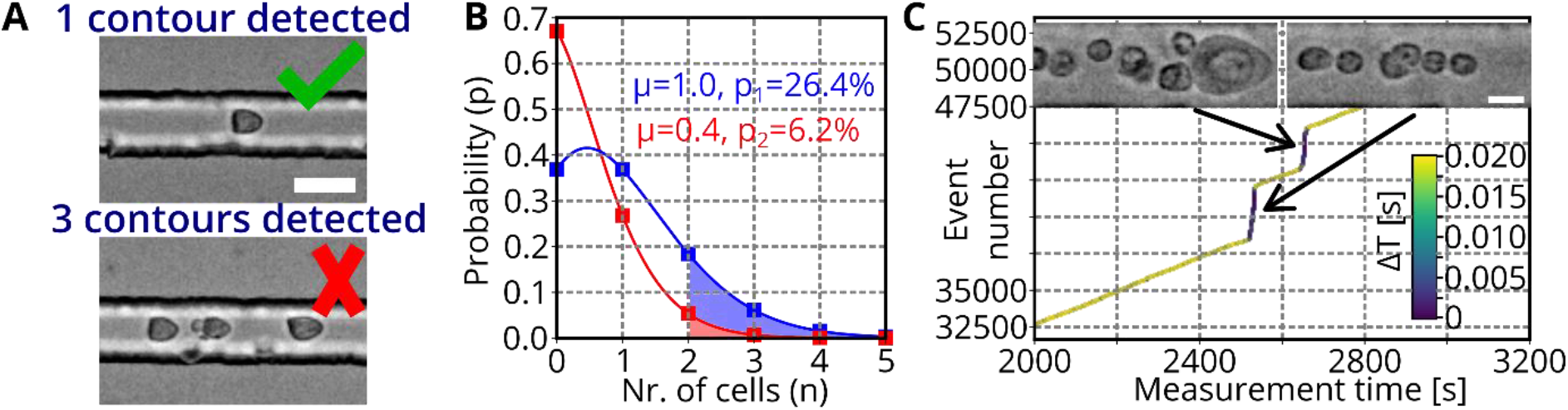
Detection and separation of cell aggregates (A) Examples of images captured during sorting. A single contour is detected in the upper image, while three contours are detected in the lower image. Sorting trigger is omitted when more than one contour is detected. Scale bar: 20 μm. (B) The histogram shows the probability to have n cells in a unit volume. The chance of having more than one cell in the sorting region during a sorting pulse is 26.4% (red) and 6.2% (blue) for an initial cell concentration of 50 million cells/ml and 20 million cells/ml, respectively. (C) Plot shows the measurement time and number of captured events of a measurement of Nrl-eGFP mice retina cells. Color code indicates the time difference between two events. While most of the time, events are captured with a time difference of >0.02 s, during an avalanche, each captured frame contains cells, resulting in a time difference of approximately 0.00033 s=0.33 ms. Scale bar: 10 μm.

The duration of a standing surface acoustic wave (SSAW) pulse is 2 ms. No additional cell should enter the SSAW region during that time to avoid accidental sorting of wrong cells. For the common flowrate of 0.04 μl/s, a volume of 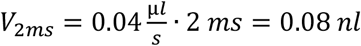 is passing the chip during an SSAW pulse. One cell contained in *V*_2*ms*_ corresponds to a concentration of 12.5 million cells/ml. To reach that concentration, an initial sample concentration of c_1_=50 million cells/ml has to be applied since the sample flow (*Q*_*sample*_ = 0.01 μ*l*/*s*) is diluted by the sheath fluid (*Q*_*sheath*_ = 0.03 μ*l*/*s*). As a result, *V*_2*ms*_ contains on average a single cell, but presence of a cell in a volume element is a random process and presence of individual cells is independent. Therefore, the number of cells (*n*) in a volume element (*V*_2*ms*_) can be described by a Poisson distribution:

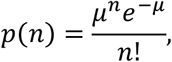

where *μ* is the expected (average) number of cells in the volume element *V*_2*ms*_. Figure 3B shows the Poisson distribution for *μ* = 1 (blue, corresponds to c_1_=50 million cells/ml). The area under the curve (pale blue) shows the probability that more than one cell is contained in *V*_2*ms*_ which is p_1_=26.4%. For sorting experiments, we reduced the concentration to c_2_=20 million cells/ml, which corresponds to an average of *μ* = 0.4 cells and a probability of getting multiple cells in *V*_2*ms*_ of p_2_=6.2% (see red plot in Figure 3B and pale red area under the curve).

The underlying assumption of the Poisson distribution is that cells travel independently, which is not entirely true, as they can stick together and form aggregates ^[17]^. As a result, avalanches of cells occasionally traverse the channel (see Figure 3C). Figure 3C shows the measurement time versus event number and the color code indicates the time difference between two captured events. Two steep increases of the curve indicate occasions where avalanches of cells flushed through the channel. As a result, each captured image contained an object, resulting in an average time difference of 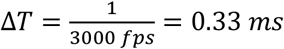 (purple regions in the plot). For the rest of the plot, the event number rises steadily and the time difference between captured events is on average 0.09 s (yellow regions of the line), which is a bit lower than the expected frequency. This is likely caused by cell sedimentation over time. Figure 3C suggests that avalanches of cells can be identified based on the characteristic time difference between captured events of 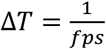. Therefore, we implemented a timer, allowing to suppress the sorting pulse if Δ*T* is below a set threshold. In practice, we found that a Δ*T* of 0.38 ms results in reliable omission of sorting during cell avalanches. The image insets in Figure 3C deliberately show only events with multiple cells in an image. While such events occur more often during avalanches, the majority of the images still shows a single cell. This fact highlights the advantage of time delay analysis in contrast to contour count. All methods were implemented into the C++ based sorting software.

### DNN architecture for optimized CPU utilization

Intelligent image-activated cell sorting allows to sort cells based on the decision of a trained DNN. While a CNN would be the preferred architecture for image classification tasks, in ref. ^[11]^, a multilayer perceptron (MLP) was used due to considerably better computational efficiency. The input layer of the model accepts grayscale values of an 8-bit raw image divided by 255. The following hidden layers perform a transformation of the input information by a set of weights and biases and an activation function (Rectified linear unit – ReLU) as indicated in Figure 4A. The MLP was trained to return the class probabilities for different blood cell types in the output layer. To allow for real-time inference to trigger a cell sorting mechanism, the MLP was optimized to provide an inference time *t* < 200 μ*s* while conserving a high classification accuracy for the distinction between different blood cell types. However, CPU specifications were not regarded in the choice of MLP design. Modern CPU chipsets provide methods for parallel computations (Hyper-Threading, Intel Advanced Vector Extensions), allowing to increase the complexity of an MLP, without changing its inference time.

**Figure 4:**
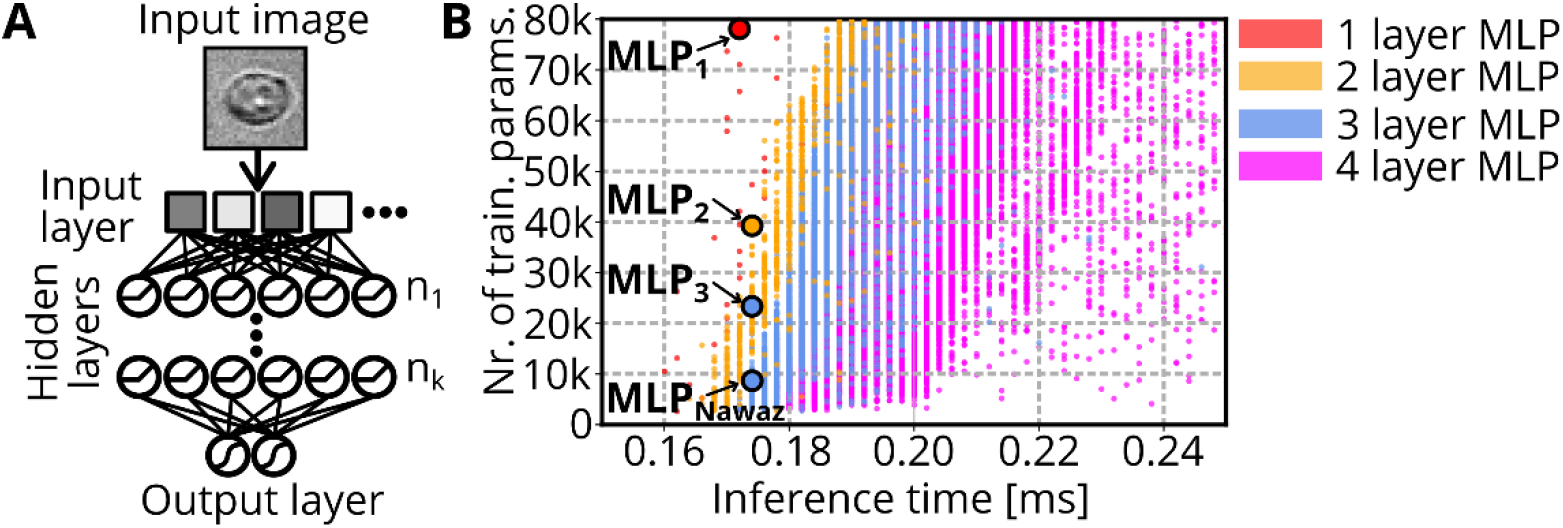
MLP screening (A) Sketch shows general design of multilayer perceptrons. The input layer contains all pixels of the provided image. Each of the following *k* hidden layers contains *n*_*i*_ (1 ≤ *i* ≤ *k*) nodes. Each node represents a linear combination of the input values, which is modulated by an activation function (ReLU for the hidden layers and Softmax for the output layer). The output layer returns probabilities for each class of the classification task. (B) The scatterplot shows the inference time and number of trainable parameters of 396,521 different MLP architectures with *k*=1 (red), *k*=2 (orange), *k*=3 (blue), *k*=4 (magenta). Chosen models with identical inference time, but more trainable parameters compared to MLP_Nawaz_ are indicated by MLP_1_, MLP_2_, and MLP_3_.

The complexity of an MLP depends on its number of parameters. The number of parameters increases the more layers and nodes are present in the neural net. Therefore, we built MLPs with *k* (1 ≤ *k* ≤ 4) hidden layers and iterated through a set of numbers of nodes *n*_*i*_ (Figure 4A). The number of nodes *n*_*i*_ of each layer was set to a multiple of 8 between 8 and 240 and for every possible combination, a model was built to determine the inference time and the number of trainable parameters *N*. To limit the computational resources, we omitted models containing *N* >80,000 parameters from the screening, resulting in a total number of 396,521 models (30, 671, 16527, and 379,293 models for *k*=1,2,3, and 4, respectively). The screening was carried out on the same PC that is used to operate the soRT-FDC setup (Intel Core i7 3930K @ 3.2GHz) and results are shown in Figure 4B (red, orange, blue, and magenta indicate models for *k* =1,2,3, and 4, respectively).

As expected, MLPs with more layers but the same number of parameters have a higher inference time due to reduced potential of parallel computation. The MLP architecture suggested by Nawaz et al. ^[11]^ which has *N*=8708, is included in our screening, and results in an inference time of *t*_*Nawaz*_ = 174 μ*s* (indicated as MLP_Nawaz_ in Figure 4B). Interestingly, no 4-layer MLP reached an inference time ≤ *t*_*Nawaz*_. Multiple models with k=1,2, and 3 comprehend more trainable parameters while having an inference time close to *t*_*Nawaz*_. We searched for models with the maximum number of parameters in the range 170 μ*s* ≤ *t* ≤ 175 μ*s*. The identified models with *k*=1,2, and 3 layers are indicated in Figure 4B by MLP_1_, MLP_2_, and MLP_3_, respectively. The models MLP_1_, MLP_2_, and MLP_3_ contain 2.7 to 9.1 times more trainable parameters compared to MLP_Nawaz_ and the total number of parameters for each model is shown in Table 1. The screening is independent of actual classification performance, but allows to find models with optimized CPU utilization. In the following, these models are employed to solve an image classification problem to assess the resulting accuracy levels.

**Table 1:**
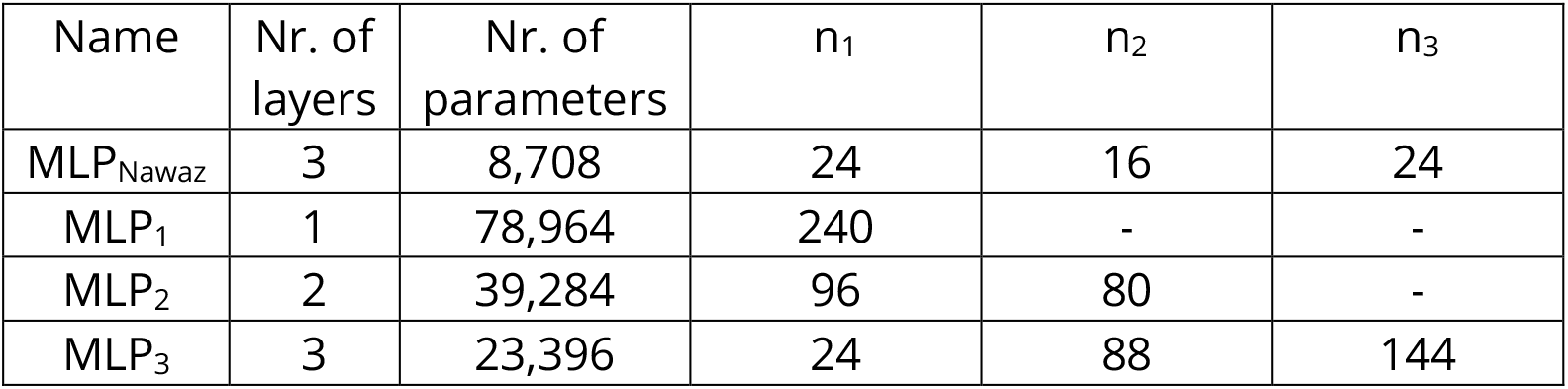
MLP screening summary

| Name | Nr. of layers | Nr. of parameters | $n_1$ | $n_2$ | $n_3$ |
| --- | --- | --- | --- | --- | --- |
| MLP<sub>Nawaz</sub> | 3 | 8,708 | 24 | 16 | 24 |
| MLP<sub>1</sub> | 1 | 78,964 | 240 | - | - |
| MLP<sub>2</sub> | 2 | 39,284 | 96 | 80 | - |
| MLP<sub>3</sub> | 3 | 23,396 | 24 | 88 | 144 |

### DNN classifier for photoreceptor detection and sorting

We performed seven independent experiments using RT-FDC to acquire data from dissociated retinae of Nrl-eGFP mice at postnatal day 4 (P04)±1 day. To that end, we used the Nrl-eGFP mouse line, which expresses eGFP under the control of the Nrl promoter, labelling rod photoreceptors from an early stage onwards. Figure 5A shows an example measurement and gates indicate certain subpopulations of cells. In a size region between 20 and 35 μm^2^, there are cells of various fluorescence expressions. To minimize wrongly labelled cells in the dataset, we employed CNN_doublet_ to remove all events with p_doublet_>0.3, excluding doublets and too proximate cells. Furthermore, we used a conservative gating strategy by only keeping cells with very low and very high fluorescence for the class of small GFP^−^ and small GFP^+^ cells, respectively (see gray and green rectangles in Figure 5A). Debris (area<20 μm^2^) and objects larger than 35 μm^2^ were not considered for the deep learning image classification task as they can be gated out based on their size during sorting. The challenging classification task that should be solved using DNNs is to distinguish small GFP^+^ (green in Figure 5A) and small GFP^−^ cells (gray in Figure 5A).

**Figure 5:**
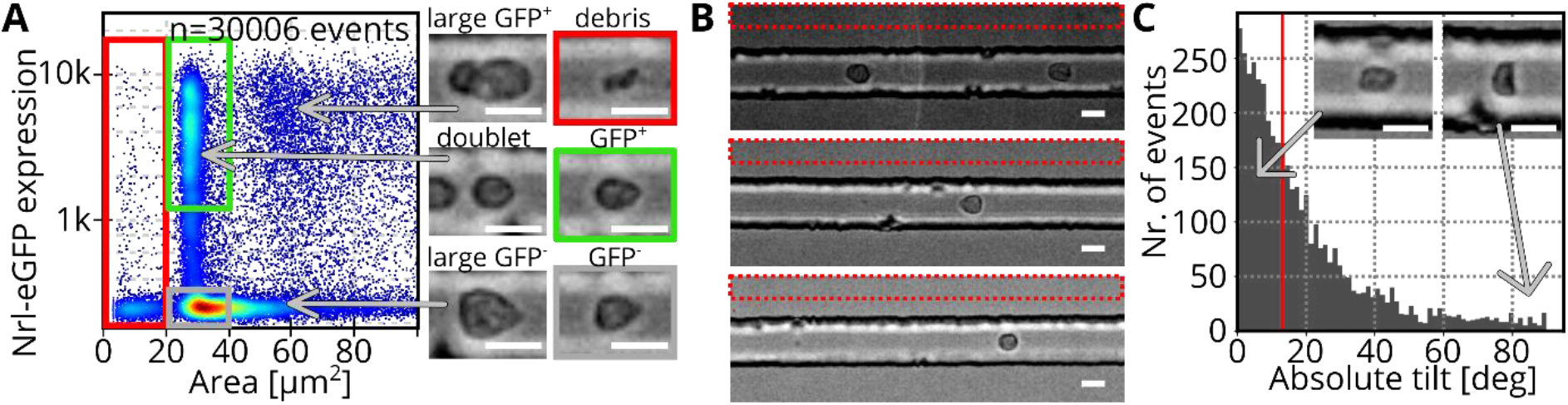
Dataset assembly (A) The scatterplot shows a measurement of dissociated retina (Nrl-eGFP) in soRT-FDC. Axes show the cell size (area in μm^2^) and the fluorescence expression of Nrl-eGFP. Red, green and gray rectangles indicate regions in the plot which correspond to debris, small GFP^+^, and small GFP^−^ cells, respectively. Images show examples of the appearance of cells at different locations in the scatterplot. The color code indicates the density of data points. Scale bars: 10 μm. (B) Images show three different measurements with various brightness levels. To evaluate the background brightness and image noise, a region above the cannel was used (red rectangle). Scale bars: 10 μm. (C) Histogram shows the absolute tilt of contours of small GFP^+^ events (same measurement as shown in Figure 5A). The red line indicates the median tilt at 13°. Image insets show exemplary phenotypes of cells at low (left) and high (right) tilt. While a low tilt indicates a good alignment with the flow, a tilt of 90° shows a cell aligned orthogonal to the flow direction. Scale bars: 10 μm.

In the current experimental setup, the focus is adjusted manually, resulting in slight differences between sessions and even slight focus drifts during long sorting procedures. To include phenotypes from different focus positions in the dataset, the focus was manually altered during acquisition of the training dataset. The range of alteration was kept in a range that would in practice be used for sorting or measurement. For acquisition of the validation dataset, the focus was left at a fixed position. Table 2 shows the number of events captured for small GFP^−^ and small GFP^+^ cells.

**Table 2:** Number of images in training and validation set.

| Class name | Nr. of training images | Nr. of validation images |
| --- | --- | --- |
| Small GFP<sup>-</sup> | 52,127 | 14,000 |
| Small GFP<sup>+</sup> | 43,321 | 14,000 |

As focus alteration increases the variety of phenotypes contained in the training dataset we would like to introduce the phrase “experimental data augmentation”. In contrast, “mathematical data augmentation” refers to computational operations applied to the image data after the measurement. Mathematical data augmentation allows to modify the image phenotype during DNN training and was shown be an effective tool to improve the accuracy and robustness of DNNs ^[18]^. A strong modification of the phenotype may enable the DNN to become robust to such alterations, but also increases the difficulty to converge. Therefore, data augmentation should ideally modify the images in a range that could occur in practice. In the following, image augmentation operations are introduced and assessed to identify sensible parameter settings. Each augmentation option is implemented into AIDeveloper which is a software for training DNNs for image classification without need for programming. All model training in this study has been performed using AIDeveloper 0.2.3 ^[19]^.

#### Data augmentation: Brightness

In the current soRT-FDC setup, there is variation in brightness between experiments. Alteration of brightness can be performed computationally. To get an intuition for the range of brightness levels of different experiments, we assessed pixels at the upper border (10×255 pixels, see red rectangles in Figure 5B) of one image from each of 29 measurements. This region allows to obtain information of the background brightness as it is located outside the measurement channel. We found minimum, maximum, and median brightness levels of 24, 49, and 39, respectively. For each case, one example image is shown in Figure 5B and a histogram of the background brightness values of 29 measurements is shown in Figure S4 A, Supporting Information. AIDeveloper allows a linear brightness alteration: *I*′ = *mI* + *n*, where *I* is the original image and *m* and *n* are random values. Assuming an image of median brightness level, the full range of brightness levels could be covered using a multiplicative factor of 0.6 < *m* < 1.3 (given *n* = 0). Similarly, the range of brightness values could approximately be covered using *n* = ±12 (given *m* = 1).

#### Data augmentation: Gaussian noise

The captured images of soRT-FDC already contain image noise, which is static, i.e. the same noise pattern is present in each image. By applying Gaussian noise, the noise pattern of an image can be altered. To determine the level of noise in original images, the standard deviation of the pixel values in the background regions (red rectangle in Figure 5B) was determined for each measurement of the training and validation set. In average, the standard deviation of the background pixels is 2.9. For a histogram of the values from 29 experiments, see Figure S4 B, Supporting Information.

#### Data augmentation: Rotation

In the current soRT-FDC setup, there is variation of the alignment of individual cells in the channel. Rotation of images can also be performed computationally. To assess the typical range of rotational variation of cells, the tilt of the contours was determined for small GFP^+^ cells of all measurements of the training and validation set. Tilt is computed based on the contour of the tracked object (see Materials and Methods). The tilt of the small GFP^+^ events of the measurement displayed in Figure 5A is shown in Figure 5C. The red line indicates the median tilt, located at 13°, meaning, 50% of cells have a tilt ≤13°. In average, the median tilt across all measurements of the training and validation set is 11°.

#### Data augmentation: Flipping

The measurement principle of RT-DC infers a vertical symmetry, allowing to perform a vertical flipping of images without changing the typical phenotype. In contrast, horizontal flipping would result in unusual phenotypes as cells are deformed in the channel according to the flow direction. Therefore, only vertical flipping is a useful image augmentation operation for RT-DC, RT-FDC and soRT-FDC data.

#### Data augmentation: Left-right, up-down shift

During a measurement or during sorting, the contour of each object is tracked in real-time and the bounding box is determined. To crop the original image of size 80×250 down to 18×18 pixels with the cell body centered, the middle of the bounding box is used. Image noise affects the location of the contour and the resulting middle point of the cell. Therefore, we used random shifting (left-right and up-down) of the cropped image by one pixel during model training.

#### Learning rate screening

The error of a model is determined by a loss function *L* (categorical cross entropy) and stochastic gradient descent allows to find in which direction the weights of a model (*W*) need to be updated in order to reduce the loss 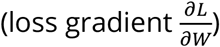 ^[20]^. The learning rate (*l*) is one of the most important hyper-parameters when training DNNs as it controls how strong the weights *W* of a model are adjusted in each training iteration (*n*):

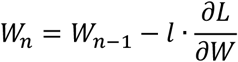

The framework for deep learning which was used in this study (TensorFlow, Keras) suggests a default learning rate of *l* = 0.001, but there is no guarantee that this is an optimal value for each dataset and model ^[21,22]^. Low learning rates correspond to slow learning of the model and therefore unnecessarily long training times. On the other hand, high learning rates can result in strong weight updates, preventing from reaching the minimum, or even a divergence of *L*. To discover a sensible value for *l*, a screening of a range of learning rates can be performed ^[23]^. To provide an easy access to that method, we implemented it into AIDeveloper. Graphical software elements guide the user through the analysis and tooltip annotations offer basic information (see Figure S5, Supporting Information). To our knowledge, this is the first time, the learning rate screening method is implemented into a software with graphical user-interface for easy accessibility.

#### MLP training

During acquisition of the training and validation dataset, the number of available cells was differing between samples. Therefore, some measurements contained more events than others. To avoid overfitting of the model to the phenotype of the measurement with most events, we performed random sampling to achieve an equal contribution of each measurement. In each training iteration of the model, a different batch of training images was sampled from each measurement. Using the same routine, the validation dataset was assembled before the first training iteration and left constant throughout all training iterations.

Training and validation data were loaded into AIDeveloper and the following data augmentation parameters were set: rotation: ±10°, left-right shift: ±1 pixel, up-down shift: ±1 pixel, additive brightness: ± 12, multiplicative brightness: 0.6…1.3, standard deviation of Gaussian noise: 3.0, and random vertical flipping. A learning rate screening was performed (see Figure 6A), considering the image augmentation parameters. For all MLP models, we found a steep decrease of the loss approximately at *l* = 10^−5^, which is 100 times smaller than the default learning rate (*l* = 10^−3^) as shown in Figure 6A and Figure S5 B, Supporting Information. Using the learning rate *l* = 10^−5^, the models MLP_Nawaz_, MLP_1_, MLP_2_, and MLP_3_ were trained for 30,000 training iterations (see Figure 6B, Figure S6 F, Supporting Information). Interestingly, the worst performance is shown by MLP_Nawaz_. MLP_2_ performs better than MLP_1_ despite having less trainable parameters (orange and red in Figure 6B). Table 3 shows the maximum validation accuracy for MLP_1_, MLP_2_, MLP_3_, and MLP_Nawaz_, indicating that the architecture of MLP_2_ is the best choice for this classification task. To obtain a benchmark for the classification accuracy if there was no restriction of the inference time, we trained two different convolutional neural net architectures. These architectures contain two (CNN_LeNet_) and four (CNN_Nitta_) convolutional layers (see Figure S6 D, E, Supporting Information). Interestingly, CNN_LeNet_ performs worse compared to all MLPs (see Figure S6 F, Supporting Information). Only CNN_Nitta_ was able to outperform the MLPs. For comparison, we also trained each model using the default learning rate (10^−3^) but the overall performance was lower for each model (see Figure S6 F).

**Figure 6:**
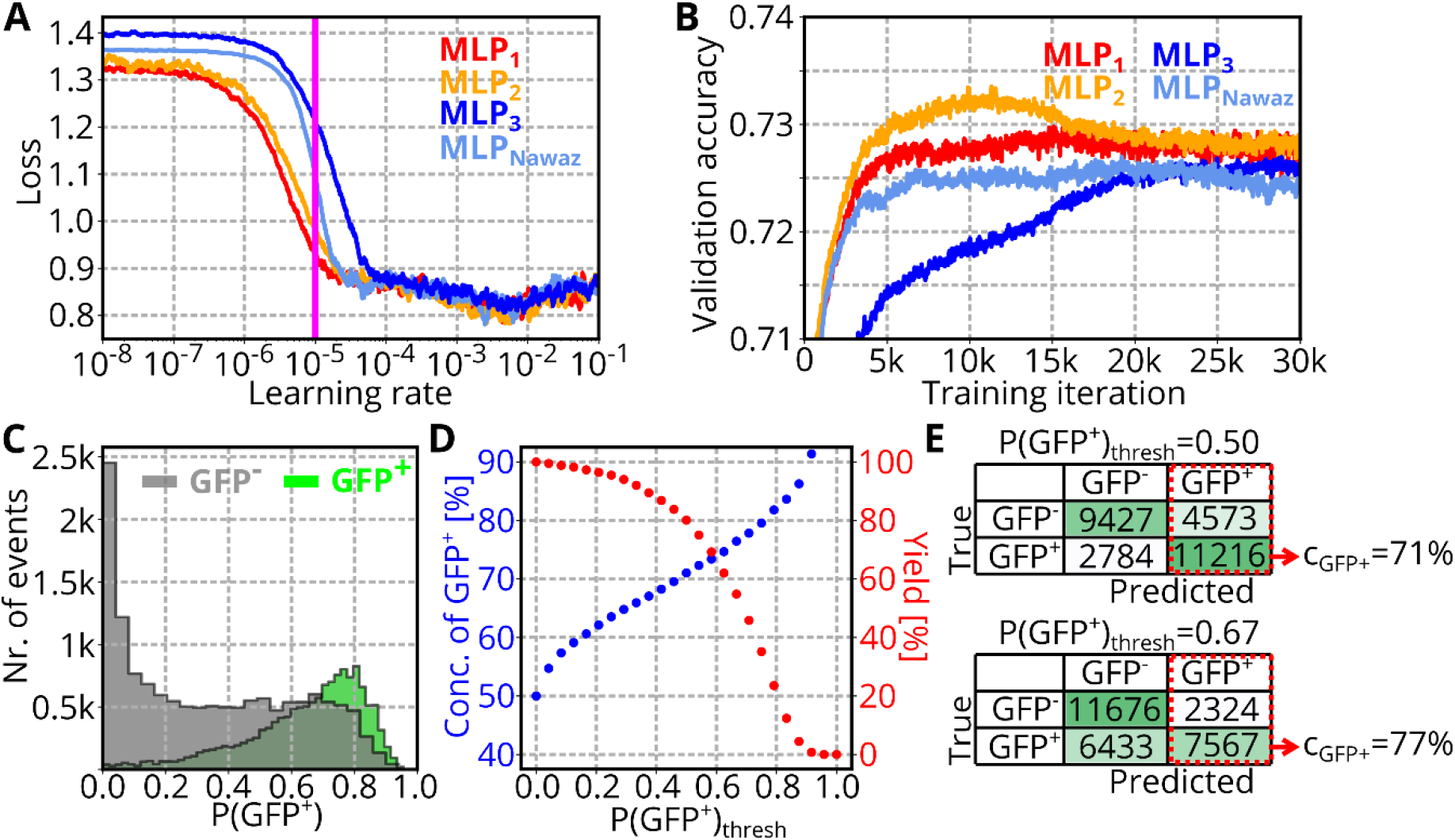
MLP training and assessment (A) Plot shows a learning rate screening for all MLP architectures. During screening, MLPs are trained using the available training data and data augmentation methods are applied. The learning rate screening was performed using AIDeveloper 0.2.3. (B) Plot shows the validation accuracy during training of four MLPs to distinguish GFP^−^ and GFP+ cells. For a smooth appearance, each line shows the rolling median (window size = 50). (C) Green and gray histogram show the probabilities returned by MLP_2_ for each event the GFP^+^ and GFP^−^ class of the validation set. (D) Scatterplot shows the concentration and yield of GFP^+^ rod photoreceptors when applying MLP_2_ to the validation set using different threshold values P(GFP^+^)_thresh_ for prediction. (E) Confusion matrices when using a threshold P(GFP^+^)_thresh_ of 0.5 and 0.67. The red rectangle indicates the events that are predicted to be GFP^+^. Those events would be sorted during a sorting experiment.

**Table 3:** Comparison of best models based on the maximum validation accuracy (max. val. acc.) of each MLP architecture.

| Name | Max. val. acc. | Training iterations to reach max. val. acc. | Training time [h] to reach max. val. acc. |
| --- | --- | --- | --- |
| MLP<sub>1</sub> | 0.735 | 21,790 | 37.3 |
| MLP<sub>2</sub> | 0.737 | 11,442 | 20.1 |
| MLP<sub>3</sub> | 0.732 | 21,271 | 35.5 |
| MLP<sub>Nawaz</sub> | 0.731 | 25,024 | 38.5 |

When applying MLP_2_ to an image, the model returns the probability that the image contains a small GFP^+^ cell: P(GFP^+^). The histogram in Figure 6C shows P(GFP^+^) for all events of the validation set. As expected, events that are actually GFP^+^ cells return high values for P(GFP^+^) (green histogram), while GFP^−^ cells tend to return lower P(GFP^+^) values (gray histogram). But there is also a considerable overlap between the distributions, which is the reason for the imperfect classification performance of the model. Typically, a threshold of P(GFP^+^)_thresh_=0.5 is used to assign events to different classes. By increasing this threshold, only cells are predicted to be GFP^+^ where the model returns a high enough P(GFP^+^). Increasing P(GFP^+^) causes an increase of the precision (see Materials and Methods), which would in practice correspond to a higher concentration of GFP^+^ cells in the target sample after sorting:

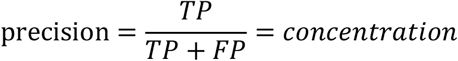

(with TP: true GFP^+^, FP: false GFP^+^). At the same time, increasing the threshold reduces the sensitivity of the model, which in practice means a reduced yield of GFP^+^ cells after sorting:

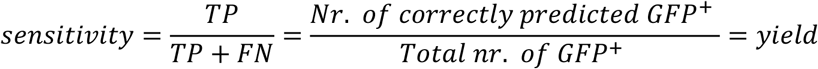

(with FN: false GFP^−^). The evolution of concentration and yield for different threshold values is plotted in Figure 6D.

For one photoreceptor transplantation experiment, 100,000 cells are required and the sorting duration should be limited to one hour to assure high viability of the cells ^[24]^. Calculations above showed that in average 0.4 cells are passing the camera within 2 ms (for a sample concentration of 20 million cells/ml). As a result, in average one cell is captured every 5 ms, which corresponds to a measurement frequency of 200 cells/s. As there are approximately 50% GFP^+^ cells, 100 cells/s could potentially be sorted. Due to the presence of cell aggregates, a more realistic sorting rate is 75 cells/s. Based on these boundary conditions, the minimum yield can be computed as following:

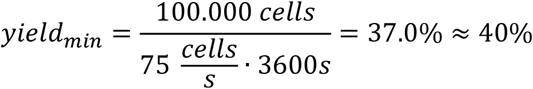

The yield of 40% is reached for a P(GFP^+^)_thresh_ of 0.67 (marked in plot), which corresponds to a concentration of GFP^+^ cells of 77%. Figure 6E shows confusion matrices for P(GFP^+^)_thresh_=0.5 and P(GFP^+^)_thresh_=0.67.

### Photoreceptor sorting and transplantation

To verify the working principle, we employed the methods introduced in this work for image-based sorting of rod photoreceptors of dissociated Nrl-eGFP mouse retina. After sorting, the initial sample and the sorted target sample were both measured using RT-FDC to evaluate the number of fluorescent cells. The color code of scatter plots in Figure 7 illustrates the event-density, which suggests that the maximum density is located at 300 and 4,000 a.U. of fluorescence for the initial and target sample, respectively. An elevated fluorescence of cells in the target sample is also confirmed by the medians of the fluorescence intensity (M_Init_=728 and M_Targ_=1,684 in Figure 7A). To evaluate the number of GFP^+^ and GFP^−^ events a gate was chosen manually (solid green line in Figure 7A). The percentage of events within that gate is 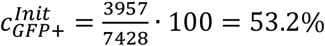 for the initial sample and 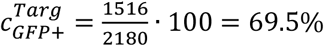 for the target sample.

**Figure 7:**
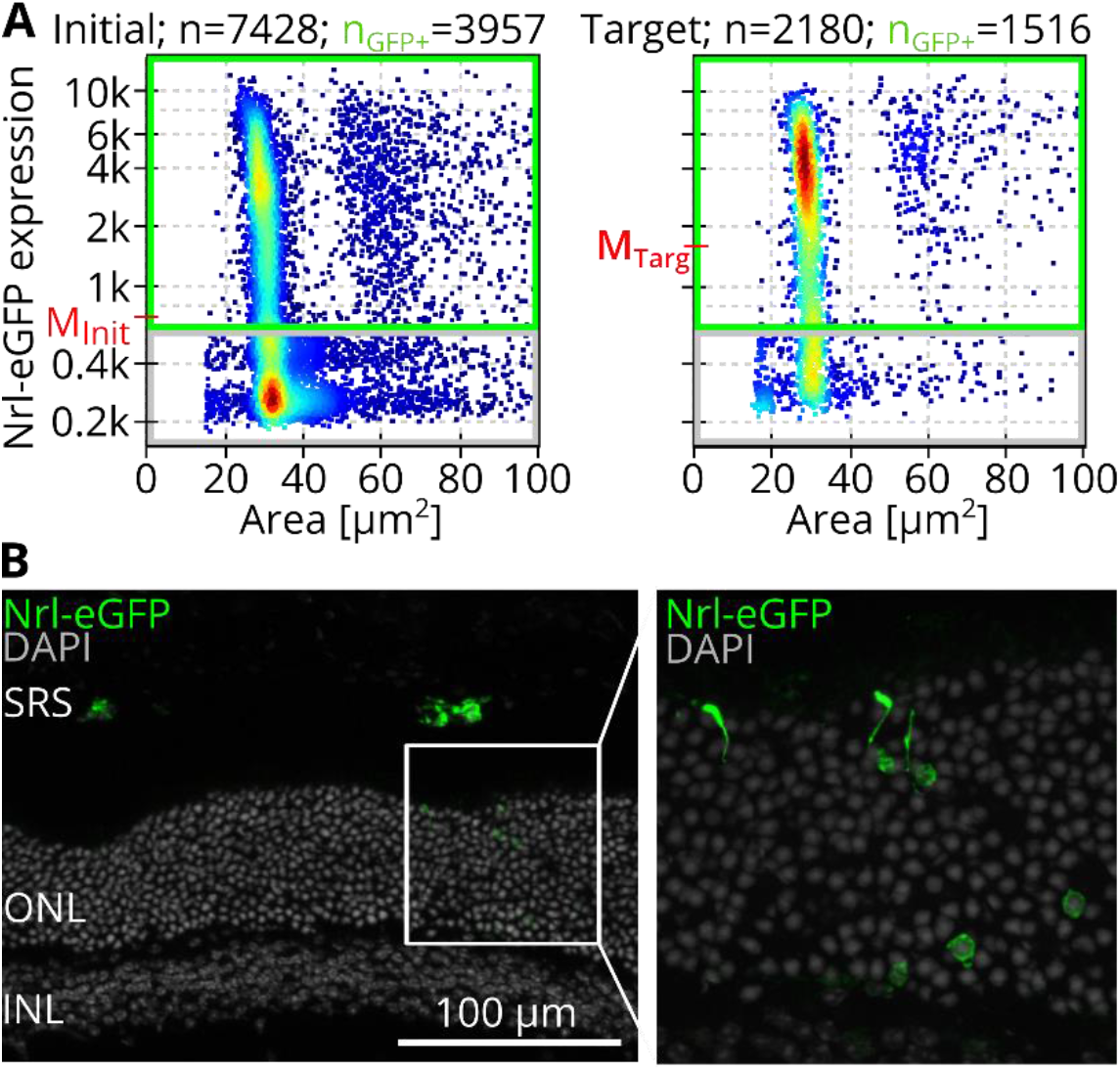
Photoreceptor sorting of dissociated Nrl-eGFP mouse retina cells & transplantation (A) Scatterplots show RT-FDC measurements of the initial sample and the target sample after sorting. The axes show the area and fluorescence expression and the color code represents the density of events. The median fluorescence expressions are given as M_Init_ (= 728) and M_Targ_ (= 1,684). The gating strategy for selection of GFP^+^ events is indicated by a green rectangle, resulting in 53.2% and 69.5% GFP^+^ cells in the initial and target sample, respectively. (B) Immunofluorescence images showing sorted GFP^+^ cells in the murine SRS, two weeks after transplantation. GFP^+^ cell bodies and segments can be found in the host ONL (magnification), likely as a result of cytoplasmic material transfer from donor to host cells. SRS = subretinal space; ONL = outer nuclear layer; INL = inner nuclear layer.

Cells contained in the target fraction were washed and subretinally transplanted into adult female C57Bl/6JRj mice. Two weeks after transplantation, GFP^+^ signal could be detected marking transplanted cells in the subretinal space of recipient mice (Figure 7B), as well as in photoreceptor cell bodies within the host ONL (Figure 7B, insert), the latter likely as a result of material transfer from donor to host cells ^[25]^. Although control eyes, in which similar numbers of unsorted cells were transplanted, contain more GFP^+^ cells at analysis (Figure S9, Supplementary Information), this is a proof of concept that cells enriched via soRT-FDC can be used for transplantation and survive in the murine retina, making soRT-FDC a useful method to provide cells for downstream applications.

## Discussion

High-throughput imaging flow cytometry with the option for label-free cell classification and sorting has the potential to complement biomedical research as it removes the dependency on molecular labels. In this work we introduce methods to improve reliability of soRT-FDC with respect to data analysis and cell sorting of dissociated tissues.

### Hardware based reduction of cell aggregates

As soRT-FDC is based on a measurement of single cells in a narrow channel, we introduce a microfluidic chip design that promotes division of cell aggregates by means of serpentines and filter structures (see Figure 1C). The majority of cells contained in human retinal organoids, young postnatal mouse retina, and human blood have a diameter below 15 μm. For the sample inflow, the minimum distance of the filter structures is therefore 15 μm, allowing most cells to pass without mechanical impact. A better separation of aggregates could be achieved using filters with an even smaller distance, or higher flowrates. We also placed filter structures in the sheath inflow part with an even smaller minimum inner distance of 10 μm. As pre-clinical experiments are typically not carried out in a clean-room atmosphere, this filter structure serves to catch dust particles and also debris of the PDMS chips, which occur even in the sheath inlet and the filter structures allow to catch those objects.

### DNN based detection of cell aggregates

While doublet detection is not trivial in FACS as doublets are defined only indirectly through the ratio of event size and width, in imaging flow cytometers like soRT-FDC, a bright-field image is available for visual inspection ^[26,27]^. Because of the large datasets, containing several thousands of images, an automated doublet detection is desired. Many doublets can be identified by a larger area or less smooth contour. Interestingly, many HRO photoreceptors show a characteristic shape with an appended tail (Figure 2A), likely presenting neural processes such as inner and outer segments. With such irregular morphologies, area and area ratio (measure for smoothness of contour) are insufficient to distinguish cell doublets from singlets. Since single cells and doublets of cells can be categorized by the human eye, we manually trained a convolutional neural net for doublet detection using data from HROs. When creating the dataset, events were also labeled as doublet when a second object travelled too closely (distance <15 μm) for confident assignment of the fluorescence intensities to the correct cell (see Figure S3 A, Supporting Information). Due to its reliable capacity of avoiding wrong assignment of fluorescence signals in RT-FDC datasets and detecting aggregates, this model will also be helpful for RT-DC datasets where cells possess a heterogeneous shape and are likely to cluster, e.g. when assessing activated neutrophils ^[28]^.

#### Labelling software: YouLabel

To facilitate the manual labelling process, we developed YouLabel. YouLabel is a software with graphical user-interface allowing to view images of an RT-DC, RT-FDC or soRT-FDC dataset and perform binary labelling. YouLabel is especially useful to screen large datasets for rare events such as doublets. The open-source software is provided as an executable for Windows and as Python script on GitHub: https://github.com/maikherbig/YouLabel. YouLabel cannot only be employed for retina datasets or datasets of dissociated samples, but for any (RT-FDC) dataset which needs to be split into two groups. However, splitting into more than two groups is currently not supported.

Despite originating from a different species and more than one year between capturing of training and testing dataset, the doublet discrimination model shows a robust classification performance for primary mouse retina cells and even for cell types of entirely different lineages, such as blood cells (see Figure S3 C, Supporting Information).

So far, the doublet discrimination model was only trained using data from a single microscope system using fixed settings and sorting chips. As the substrate of sorting chips is birefringent, the phenotype is different from normal glass chips. Therefore, the model will likely fail to make correct predictions for images of cells in chips with glass substrate, different magnification levels or illumination. To optimize the model for altered system settings, transfer learning could be employed ^[29]^. Such an approach requires the acquisition of only a small dataset using the new system, which can then be employed to continue training the existing model. Building on pre-existing capabilities, transfer learning has the advantage of saving time and computational power compared to training a new model from the beginning.

### Detection and separation of cell aggregates for single cell sorting

Due to the computational time required, we unfortunately could not use the doublet discrimination model during sorting. To recognize cell aggregates and avalanches of cells, we instead used the number of contours and the time delay between detected objects. However, these methods are purely image based and are therefore limited to the framerate of the camera. By integrating a laser for acquisition of forward scattered light, the cell count could be tracked for up to 50,000 cells/s (similar to FACS), which would allow for improved detection of cell avalanches ^[6]^.

To further manually decrease the occurrence of cell aggregates, the cell concentration could be reduced, leading to an increase of the average space between cells in the sample and thus in the measurement channel (Figure S7 A, Supporting Information). For example, for concentrations of 50, 20, and 10 million cells/ml, the free space between cells in the channel is 190 μm, 490 μm, and 990 μm, respectively (assuming each cell has a diameter of 10 μm). However, high measurement throughput requires high cell concentrations (Figure S7 B, Supporting Information e.g. average measurement frequency of 500, 200, and 100 cells/s at 50, 20, and 10 million cells/ml, assuming a total flow rate of 0.04 μl/s), so an optimization of aggregate prevention without the need for sample dilution would be preferable.

### DNN architecture for optimized CPU utilization

The most popular DNN architecture for image classification is the CNN. In CNNs, convolutional filters are fixed and applied across the image to transform the pixels of a certain neighborhood, which reduces the degrees of freedom of the model. This approach often results in a robust classification performance of CNNs, but comes at a cost of computational time, because convolutional filters are facilitated by sparse matrix operations. To run inference using CNNs with sub-millisecond inference time, special hardware such as FPGAs are required, rendering CNNs unfavorable for sorting when computation capacities are limited to a CPU ^[8]^. The requirement for low inference time calls for utilization of very small models with a low number of parameters. MLPs allow an efficient usage of the available parameters as every node in a layer is connected to each node in the subsequent layer. However, the increased flexibility of MLPs makes them more prone to overfitting. In this work, we performed a screening to identify MLPs that offer a high number of trainable parameters at low inference times. However, the result of that screening is only valid for our PC hardware and power settings. Therefore, we provide the script (Zenodo: https://doi.org/10.5281/zenodo.4738936), allowing everyone to perform the screening, for example using the Python editor that is integrated into AIDeveloper.

### DNN classifier for photoreceptor detection and sorting

To train a DNN for prospective sorting, a biologically diverse dataset containing data from multiple biological replicates needs to be used. Therefore, we acquired data of seven Nrl-eGFP mice at maturation stage postnatal day 4 (P04)±1. A control of the maturation stage is required as retinal development occurs rapidly before and within 10 days after birth and it was demonstrated that murine photoreceptors at P04 are best suited for subretinal transplantation ^[30–33]^. Variation in image phenotype cannot only be due to biological, but also technical reasons. Since the substrate of microfluidic chips for sorting is a birefringent material, the phenotype differs slightly between chips. Therefore, we used a new chip for each sample. To reduce the differences between chips, a more standardized and automated chip production would be beneficial. Automation would also allow to eliminate focus and brightness differences between measurements. An autofocus system would not only omit the need to record training data at many focus positions, but also simplify neural net training as fewer degrees of freedom would have to be considered by the model.

For data acquisition, RT-FDC was employed as it allows to simultaneously record a bright field image and a 1D track of fluorescence information (see Figure S3 A, Supporting Information). The section on the fluorescence track corresponding to a certain cell in the image varies due to slightly differing object velocity in the channel. Doublets of cells, or cells travelling closely, present a risk of assigning the wrong fluorescence expression value (see Figure S3 A, Supporting Information), which would lead to wrongly labeled images in the dataset. As such, GFP^+^ cells that are larger than 35 μm^2^ refer to cell doublets i.e. photoreceptor cells attached to another cell (see example image in Figure 5A). Therefore, the dataset was cleaned through application of CNN_doublet_ and through size exclusion of cell debris (< 20 μm^2^). The remaining cells in the size region between 20 and 35 μm^2^ show a continuous range of fluorescence values, as expected for the Nrl-eGFP mouse line ^[34]^, and were considered for the DNN training process.

Prior to the MLP training, we performed a learning rate screening using AIDeveloper. We found that the learning rate *l* = 10^−5^ results in a steep decrease of the loss for all MLPs and CNNs (see Figure S5 B, Supporting Information) and trained all models using that constant learning rate (Figure 6B and Figure S6F). However, there is no guarantee that this value is optimal throughout the entire training process. Therefore, we implemented learning rate schedules (exponential decay and cyclical learning rates) into AIDeveloper, but we could not find any setting that outperformed the constant learning rate ^[23]^. In order to prevent overfitting that is more common for MLPs compared to CNNs, we introduced methods to find optimal parameters for data augmentation. While the augmentation methods are already enabled in AIDeveloper, the methods to find optimal ranges for augmentation are not implemented yet. After training using these tools, the model with the highest validation accuracy theoretically allows to enrich rod photoreceptors to 71%, or 77%, depending on the sorting threshold P(GFP^+^)_thresh_ (see Figure 6E). An established sorting method using CD73 antibodies and MACS allows to obtain a concentration of photoreceptors of ≈80% ^[24]^. MLP_2_ would theoretically also allow such a concentration when using a sorting threshold P(GFP^+^)_thresh_ of 0.76 ^[24,34]^. Unfortunately, since the curves for concentration and yield are developing in opposite direction (see Figure 6D), the corresponding yield is only 31%.

In this work we focused on MLPs due to their lower inference time. However, for image classification tasks, CNNs would be preferred as they typically allow to reach higher accuracies. For comparison, we also trained two CNNs, which required considerably longer training times, despite using a GPU (Nvidia GTX 1080). While the MLP_1_, MLP_2_, MLP_3_, and MLP_Nawaz_ reached the maximum validation accuracy after 37.3 h, 20.1 h, 35.5 h, and 42.8 hours, CNN_LeNet_ and CNN_Nitta_ took 242.0 h and 93.6 h, respectively.

The GFP^−^ cells that are in the same size region as rod photoreceptors could be any other cell type of the retina. Given the almost equal cell number of GFP^+^ and GFP^−^ cells in the size range 20 μm^2^ to 35 μm^2^, those GFP^−^ cells must either be a highly abundant cell type, or a superposition of multiple cell types. After rods, the most abundant retinal cell types are bipolar and amacrine cells ^[35]^. Unfortunately, we only had access to a bipolar reporter mouse line (mGluR6-GFP) which we measured using RT-FDC. Bipolar cell size was larger than 35 μm^2^ (see Figure S8 A, Supporting Information), indicating that the Nrl-eGFP^−^ cells cannot be caused by presence of bipolar cells. Furthermore, we measured retina cells from a cone reporter line, indicating that cones tend to be larger than 35 μm^2^ (see Figure S8 B, Supporting Information). At P04, there are still retinal progenitor cells present because cell differentiation continues until P10 ^[30,36]^. However, retinal progenitor cells are larger than rod photoreceptors ^[37]^. Furthermore, despite lacking retinal progenitor cells at P10, the Nrl-eGFP^−^ population still contains cells in the area range 20 μm^2^ to 35 μm^2^ (Figure S8 C, Supporting Information) ^[37]^. Cells that could meet the sizes of the Nrl-eGFP^−^ cells are horizontal, Müller, and amacrine cells ^[35]^. While we cannot yet specify the cell types of the Nrl-eGFP^−^ population, cell numbers suggest that amacrine cells should be assessed in the future ^[30,35]^. Currently, the MLPs are trained for the binary classification task to distinguish bright-field image differences between GFP^+^ and GFP^−^ cells. As the GFP^−^ fraction is likely composed of multiple cell types, the MLP has to learn weights that suit all of the occurring phenotypes. A more detailed labelling of cell types and training using multiple classes would allow the model to assign individual weights to each class which could result in higher accuracies.

### Photoreceptor sorting and transplantation

One common reason for blindness are retinal degenerations which result in a loss or malfunctioning of photoreceptors. A promising approach to treat retinal degenerations is photoreceptor transplantation, using photoreceptors generated in vitro from human embryonic or induced pluripotent stem cells ^[16]^. Nowadays, photoreceptors are most efficiently produced in three-dimensional organoids, which often contain a complex mix of cell types and a tissue-like architecture ^[38,39]^. Although transplantation of organoid sheets retaining that architecture is possible, it is surgically complex and might introduce undesired cell types. The donor material can alternatively be dissociated in order to obtain single cells, allowing for prospective cell sorting and purification. Established sorting techniques like MACS or FACS however require molecular markers which label the target cells ^[40,41]^. Unfortunately, such markers are not always well-defined and if they are, their binding could potentially alter the cells, which should be avoided in a clinical setting. For example, labelling photoreceptors in HROs is currently still challenging due to a lack of defined surface markers ^[16]^. Given that all parts of the soRT-FDC are disposable, the system would potentially be adaptable to GMP applications.

Here, we employ soRT-FDC to enrich young murine photoreceptors in a label-free fashion for subretinal transplantation. This proof-of-concept study shows that the cells survive the sorting process well enough to not only be detected in situ two weeks later but to also interact with the host tissue, as shown by potential material transfer to host cells (Figure 7B, insert). Despite this important progress, it must be noted that eyes transplanted with the sorted fraction contain markedly fewer cells than eyes transplanted with unsorted control samples, suggesting the sorting itself to be strenuous and potentially damaging to the cells (Figure S9, Supporting Information). Loss of cell viability is an inherent problem in most cell sorting methods ^[24]^, as high pressure and shearing forces are often required. Thus, it will be necessary to improve sorting conditions further in order to obtain better viability of the sorted populations and increase the efficiency of the sorting setup. Overall, in this report we enable soRT-FDC to perform label-free image-based sorting of photoreceptor cells from dissociated retinae of Nrl-eGFP mice. To decrease falsely labeled cells in the dataset, we present a model reliably identifying cell doublets and cells travelling too closely. For photoreceptor cell identification, we train a deep neural net using complex samples of murine retina, a tissue diverse in cell phenotypes. Different experimental and mathematical data augmentation techniques exerted upon the training dataset allowed to obtain a model with high robustness. Finally, we show that the model can be applied to make sorting decisions in a true experiment and that cells sorted with the described setup can successfully be used in downstream applications.

## Materials and Methods

### Microfluidic chip with sorting mechanism

Microfluidic chips for soRT-FDC are manufactured in house. In detail, a mixture of polydimethylsiloxane (PDMS, SYLGARD 188, Dow Corning) and curing agent (10:1, w/w) is poured over a silicon wafer master with the microfluidic design (see Figure 1C). After baking at 65 °C for 60 min, the PDMS layer is removed from the master and holes for sheath inlet, sample inlet, default outlet, and target outlets are punched using a biopsy puncher (Biopsy Punch with Plunger no. 49115, size 1.5 mm, pfm medical AG). To seal the microfluidic structures on the PDMS layer, a lithium niobate substrate is covalently bound using plasma activation (50 W, 30 s; Plasma Cleaner Atto; Diener Electronic). Bonded devices were then cured for 48 h in an oven at 65 °C.

The 128° Y-cut lithium niobate (LiNbO_3_, Roditi International) substrate is equipped with two gold electrodes (interdigital transducers - IDTs) that allow to excite surface acoustic waves for sorting. Gold electrodes were deposited onto the substrate by a metal evaporation process (NANO36, Kurt J Lesker). Each IDT has 40 electrode pairs and a distance between electrode fingers of 70 μm, which results in an excitation frequency of 55.23 MHz. By simultaneously exciting both IDTs, counter-propagating surface acoustic waves are generated, resulting in a standing surface acoustic wave (SSAW) with a wavelength of 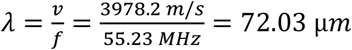 (speed of sound on LiNbO_3_ is 3978.2 m/s ^[42]^).

Objects interact with the SSAW via the acoustic radiation force ^[43]^:

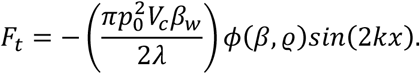

The acoustic contrast factor *ϕ*,

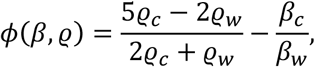

Is defined computed using acoustic pressure *p*_0_, wavelength *λ*, cell volume *V*_*c*_, cell compressibility *β*_*c*_, fluid compressibility *β*_*w*_, cell density *ϱ*_*c*_, and fluid density *ϱ*_*w*_. Since cells have a higher density (*ϱ*_*Water*_ ≈ 1.0 *g*/*m*^3^, *ϱ*_*Protein*_ ≈ 1.3 … 1.4 *kg*/*m*^3^) and a higher compressibility compared to water (*β*_*w*_ ≈ 4.5 *GPa*^−1^, *β*_*c*_ ≈ 4 *GPa*^−1^) ^[44–46]^, they have a positive *ϕ*, and therefore move towards the pressure node. The maximum translocation of an object by a SSAW is *λ*/4 = 18.01 μ*m*.

To excite the substrate, the signal of a surface acoustic wave generator (BSG F20, BelektroniG) is duplicated using two fast-switches (BPS-300, BelektroniG). LiNbO_3_ is a birefringent material. The resulting image distortion was corrected using a Polarizer (Polarizer D, Zeiss).

### soRT-FDC device

Sorting real-time deformability and fluorescence cytometry (soRT-FDC) device was assembled as shown in ^[11]^. In brief, a microfluidic chip is placed on an Inverted microscope with a 20x objective (Plan-Apochromat, 20x/0.8 NA; no. 440640-9903, Zeiss). Two syringe pumps (neMESYS 290N, neMESYS) drive sheath and sample fluid into the chip at flow rates of 0.03 μl/s and 0.01 μl/s. Another syringe pump withdraws fluid from the default outlet (−0.027 μl/s), while the target outlet is at atmospheric pressure. Within the chip, the cells flow into a narrow channel where they are aligned and slightly deformed by hydrodynamic forces. At the end of the channel, the cells are illuminated by 2 μs flashes from a high-power LED (CBT-120, Luminus Devices). The LED flashes are triggered by a high-speed camera (EoSens CL, MC1362, Mikrotron), which captures images of cells at 3000 frames per second. Image data is sent to a PC (Intel Core i7 3930K @ 3.2 GHz) via a full camera link frame grabber card (NI-1433, National Instruments) and a C++ based software analyzes images using the OpenCV library ^[47]^. A running average of the last 100 frames is computed as a background image, which is then subtracted from each subsequent image. Next, the image is binarized by a thresholding operation. In the following, erosion and dilation operations are applied to finally obtain a smooth contour from a contour finding algorithm ^[48]^. Based on the coordinates of the contour, a bounding box is computed. The middle of the bounding box is used to crop the image such that the cell body is centered. Finally, the cropped image is forwarded through a defined neural network and resulting prediction probabilities are used to trigger a sorting unit located behind the narrow channel (see Figure 1C).

Besides image acquisition, also fluorescence information can be obtained for each single cell. Fluorescence is excited using up to three lasers of wavelengths 640, 561, and 488 nm (OBIS 640 nm LX 40 mW, OBIS 561 nm LS 50 mW, OBIS 488 nm LS 60 mW, Coherent Deutschland). The laser beams form a light sheet in the middle of the region where images are captured. The emitted fluorescence signal from cells passing the light sheet is collected by a photodiode detector assembly (MiniSM10035, SensL Corporate), resulting in 1D fluorescence traces for each captured cell ^[10]^.

### Alignment of cells in the channel (tilt)

For quantification of the alignment of cells in the channel, the tracked contour is employed. The orientation *φ* of a contour with respect to 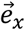 is computed by ^[49]^:

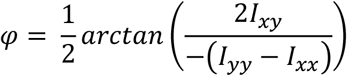

with the second moments *I*_*xx*_ = ∬_*A*_ *y*^2^*dx dy*, *I*_*yy*_ = ∬_*A*_ *x*^2^*dx dy*, and the biaxial second moment *I*_*xy*_ = − ∬_*A*_ *xy dx dy*.

### Multilayer perceptron

In an MLP, all pixel values of the image are combined using weights, biases and an activation function in a defined number of nodes. If matrix *W*^(1)^ contains the weights, *b*^(1)^ the biases and *s* the activation function of the first hidden layer, then the transformation performed by the first hidden layer can be expressed as:

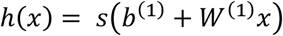

The design of layered neural networks limits the number of parallel tasks. While all nodes in a single layer could be computed in parallel, subsequent layers first require the preceding layer to complete computations. While in MLPs, each node is connected to all subsequent nodes, in convolutional neural nets the connections are limited to a certain neighborhood, defined by the size of the convolutional kernel. While this constraint typically results in more robust modes that can be trained with less data, the number of parameters, resulting in less flexibility.

### Inference time

Inference time is the time required to compute the output of an algorithm. The present work introduces different DNNs and the inference time describes how much time the PC requires to forward a single image trough the network to compute a prediction. Importantly, inference time is different from training time. During training, the model parameters are updated based on the computed gradient. The computation of the gradient is performed for batches of images in parallel. At inference time, these gradients don’t need to be computed and for sorting, only one image has to be forwarded at a time. Therefore, training of DNNs is often done on high performance computing clusters, while for single image inference, a CPU is sufficient. Sending single images to a computing cluster would result in varying data transfer times. In contrast, a local CPU can be accessed fast, especially if images are already stored on RAM.

Under low load, modern CPUs can throttle to save power, and full reactivation requires time. For exact determination of the inference time, we pre-heated the CPU by forwarding one image. Immediately after pre-heating, 500 images were forwarded through the DNN individually (not in parallel). The process of sequentially forwarding 500 images is repeated 10 times to compute an average inference time.

### Labelling software *YouLabel*

We used Python 3.5.6 to establish a software for fast labelling of events in RT-DC datasets which is open source: https://github.com/maikherbig/YouLabel. We provide a standalone executable for Windows. Alternatively, the software can be executed using the provided Python script.

### Statistical analysis

The performance of machine learning models is assessed using the confusion matrix and metrics, computed based on the confusion matrix. For computing the confusion matrix, a labelled dataset is required and the matrix shows the true and predicted label for each class. Assuming a binary classification task with a positive and a negative class, there are four options: a positive event is correctly classified as positive (true positive – TP), a positive event is falsely predicted as negative (false negative - FN), a negative event is correctly classified as negative (true negative – TN), or a negative event is falsely predicted as positive (false positive – FP). Metrics, derived from the confusion matrix are: 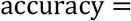 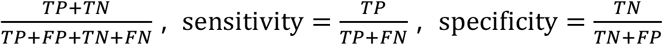 and 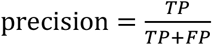. The terms “recall” and “true positive rate” are synonyms for sensitivity.

### Measurement buffer preparation

We complemented phosphate buffered saline (PBS, 10010-023, Gibco) with 10% (v/v) Leibovitz’s L15 medium (11415064, Thermo Fisher Scientific) to support viability of cells. As cells need to stay in suspension for at least one hour during sorting experiments, we added 0.6% (w/w) methyl cellulose (4,000 cPs; Alfa Aesar) to reduce sedimentation. The resulting viscosity of the buffer is 25 mPas (at 24 °C). The buffer was adjusted to pH 7.4 and an osmolality of 310–315 mOsm/kg. For reduced formation of cell aggregates, 2% (v/v) DNase stock solution was added, with DNase I stock solution containing 5 mg/ml (=10.000 Kuntz Units/ml) DNase I (DNase I, D5025-150KU, Sigma) in 0.15 M NaCl (A2942, Applichem).

### Animal welfare statement

All animal experiments were approved by the ethics committee of the Technische Universität Dresden and the Landesdirektion Dresden (approval no. TVV 10/2018 and TVV 25/2018) and performed in accordance with the regulations of the European Union, German laws (Tierschutzgesetz), the ARVO Statement for the Use of Animals in Ophthalmic and Vision Research, as well as the National Institutes of Health Guide for the care and use of laboratory animals.

### Retina single cell preparation

Neural retina leucine zipper-enhanced green fluorescent protein (Nrl-eGFP), cone-GFP and metabotropic glutamate receptor 6-GFP (mGluR6-GFP) mouse lines were used as source for rods, cones and bipolar cells, respectively ^[15,50,51]^. Single cell suspensions of P04 ±1 retina were prepared as described in ref. ^[34]^. Briefly, pups were decapitated and heads transferred to a petri dish containing cold PBS. Eyes were dissected and retinae were isolated, washed in ice-cold Cell Buffer (2 mM EDTA, 1% w/v BSA in PBS without calcium or magnesium) and transferred to 37 °C Papain solution supplied with 2.5% DNase I stock solution (> 200 KU/ml DNase I). Retinae were digested 40 min at 37 °C, with mixing of the samples by inverting the tube every 10 min. After careful manual trituration, the suspension was washed with EBSS wash (EBSS, 10% v/v DNase I stock solution, 10% v/v Ovomucoid inhibitor) after which digestion was fully stopped by overlaying onto Ovomucoid inhibitor and centrifuging for 5 min at 300 g. Supernatant was removed and cells resuspended to 20×10^6^ cells/ml in measurement buffer. Papain solution, EBSS and Ovomucoid inhibitor were taken from the Papain Dissociation System (PDS Kit, Cat. No.: LK003182, Worthington Biochemical Corporation).

### Human retinal organoid sample preparation

For analysis of human photoreceptor cells, human retinal organoids (HRO) derived from the Crx-mCherry iPSC line (kindly provided by O. Goureau, Paris) and generated as described in ref. ^[52]^ were dissociated as follows: organoids were washed 3x in 37 °C PBS, transferred to 37 °C Papain solution and incubated for 2 h at 37 °C on a horizontal shaker shaking at 90 rpm ^[53]^. Then, DNase I stock solution was added to 5% v/v (> 400 KU/ml) and HRO triturated carefully using a glass pipette. After filtering through a 30 μm filter (Cat.No.: 130-041-407, Miltenyi Biotech), cells were washed with EBSS wash, digestion was stopped and cells were resuspended as described above, centrifuging for 6 min at 600 g.

### Sample preparation for photoreceptor transplantation

For transplantation, the target, unsorted and default fractions containing 30,000 or 80,000 cells were washed with cell buffer and centrifuged for 5 min at 800 g. Cells were resuspended in transplantation (TP) buffer (Cell Buffer containing 2% v/v DNase I stock solution) transferred to a fresh tube and centrifuged again for 5 min at 800 g. The cell pellet was then resuspended in 1-2 μl TP buffer and kept on ice until subretinal transplantation. Adult (>10 w) C57Bl/6JRj females were used as recipients and trans-vitreal subretinal transplantation was performed as described in detail in ref. ^[34]^.

### Tissue processing, immunohistochemistry and imaging

Experimental animals were euthanized, eyes enucleated and fixed for 1 h in 4% paraformaldehyde (CAS: 50-00-0, Cat.No.: 100504-858, VWR)) in PBS at 4 °C. After removal of the cornea, lens, vitreous and excess muscular tissue, eyes were cryoprotected in 30% sucrose in PBS overnight and frozen in Neg-50 (Cat.No: 6502, Thermo Fisher Scientific). 12 μm sections were treated with 0.3% Triton-X100, 5% donkey serum and 1% BSA in PBS and immunostained (Primary antibody: chicken-anti-GFP, ab13970, Abcam, 1:500; Secondary antibody: donkey-anti-chicken-Cy2, 703-225-155, Jackson Immuno Research, 1:1000; Counterstain: DAPI, Sigma, 0.2 μg/ml). Stained sections were imaged using an Apotome Imager Z1 equipped with ApoTome.2 and ZEN 2.5 pro blue edition (Carl Zeiss Microscopy GmbH).

## Supporting information

Supporting Information is available from the bioRxiv preprint server or from the author.

## Data availability

All datasets and trained models are publicly available at Zenodo: https://doi.org/10.5281/zenodo.4738936. Python scripts to reproduce analysis tasks and generate plots are contained in the repository. Each script can be executed using PyBox 0.1.0 (https://github.com/maikherbig/PyBox). PyBox 0.1.0 is a readily installed Python environment that contains all required Python packages.

AIDeveloper is open source software and can be downloaded from GitHub: https://github.com/maikherbig/AIDeveloper.

YouLabel is open source software and can be downloaded from GitHub: https://github.com/maikherbig/YouLabel.

## Acknowledgement

We like to thank Olivier Goureau for providing the human retinal organoid Crx-mCherry iPSC line. The authors acknowledge funding from the Deutsche Forschungsgemeinschaft (DFG Project number 399422891 to M.A) within the SPP2127 Program (AD375/7-1 to M.A. and GU612/5-1 to J.G.). We further like to thank the animal facility staff at CMCB Dresden.

## Author contributions

M.A. and J.G. conceived the project. M.H., K.T., T.F., O.B, and S.G. performed experiments. M.H. and K.T. wrote the manuscript. M.N programmed the software controlling the sorting device. A.A.N. manufactured the microfluidic chips for sorting and contributed to the design of those.

## Competing interests

All authors declare no competing interests.

## Notes

https://doi.org/10.5281/zenodo.4738936

### Competing Interest Statement

The authors have declared no competing interest.

