## Supporting Information for "Label-free imaging flow cytometry: analysis and sorting of enzymatically dissociated tissues"

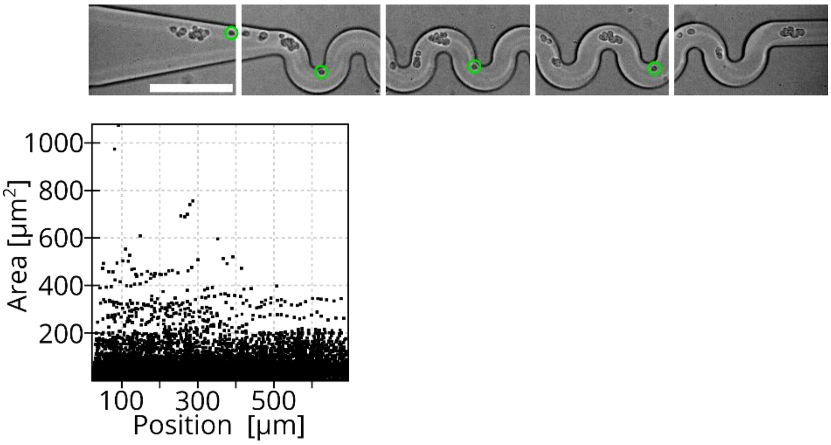


Figure S1: Reduction of aggregates in the serpentine

Top image shows multiple snapshots of one aggregate of cells travelling through the serpentine. Cells are closely together before the serpentine (left). Within the serpentine, they are separated into single cells and smaller aggregates and the spacing between cells increases. For a guide of the eye, the identical cell is marked using a green circle in each snapshot. Scale bar: 100 µm.

For a more quantitative approach, a measurement was conducted in the region of interest as indicated by the image above. The scatterplot indicates that large aggregates with sizes above 400 µm$^2$ are only present at the beginning (position 0 to 400 µm) of the serpentine.

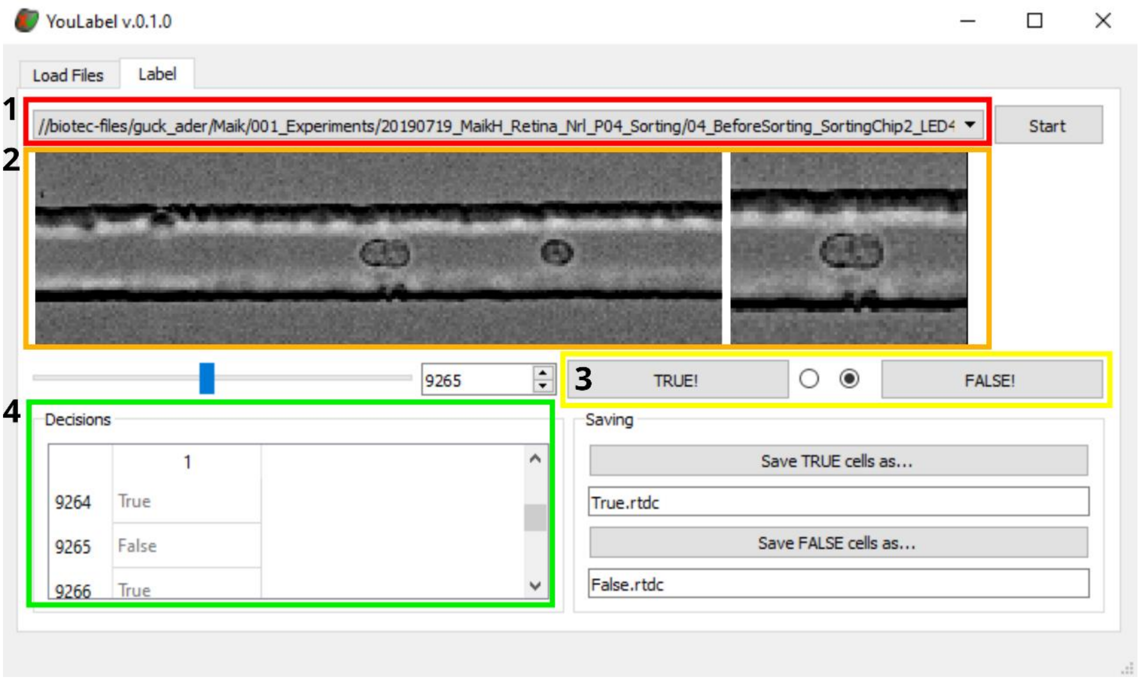


Figure S2: Screenshot of labelling software "YouLabel"
Colored rectangles in the image indicate the workflow
1 – Select dataset that should be worked on. After selection the file is loaded and the images of the dataset are displayed.
2 – Images of the loaded dataset are displayed. In RT-FDC datasets, the midpoint of each tracked object is stored. Based on this midpoint, the full image (left) is cropped (right) to show the tracked object in the center. To display the next or the previous image, the right and left arrow key can be used.
3 – Buttons to indicate if the displayed image corresponds to class 1 (True) or class 2 (False). Alternatively, the keyboard shortcuts "T" and "F" may be used.
4 – List shows the labelling decisions. By default, all events are labelled "True".

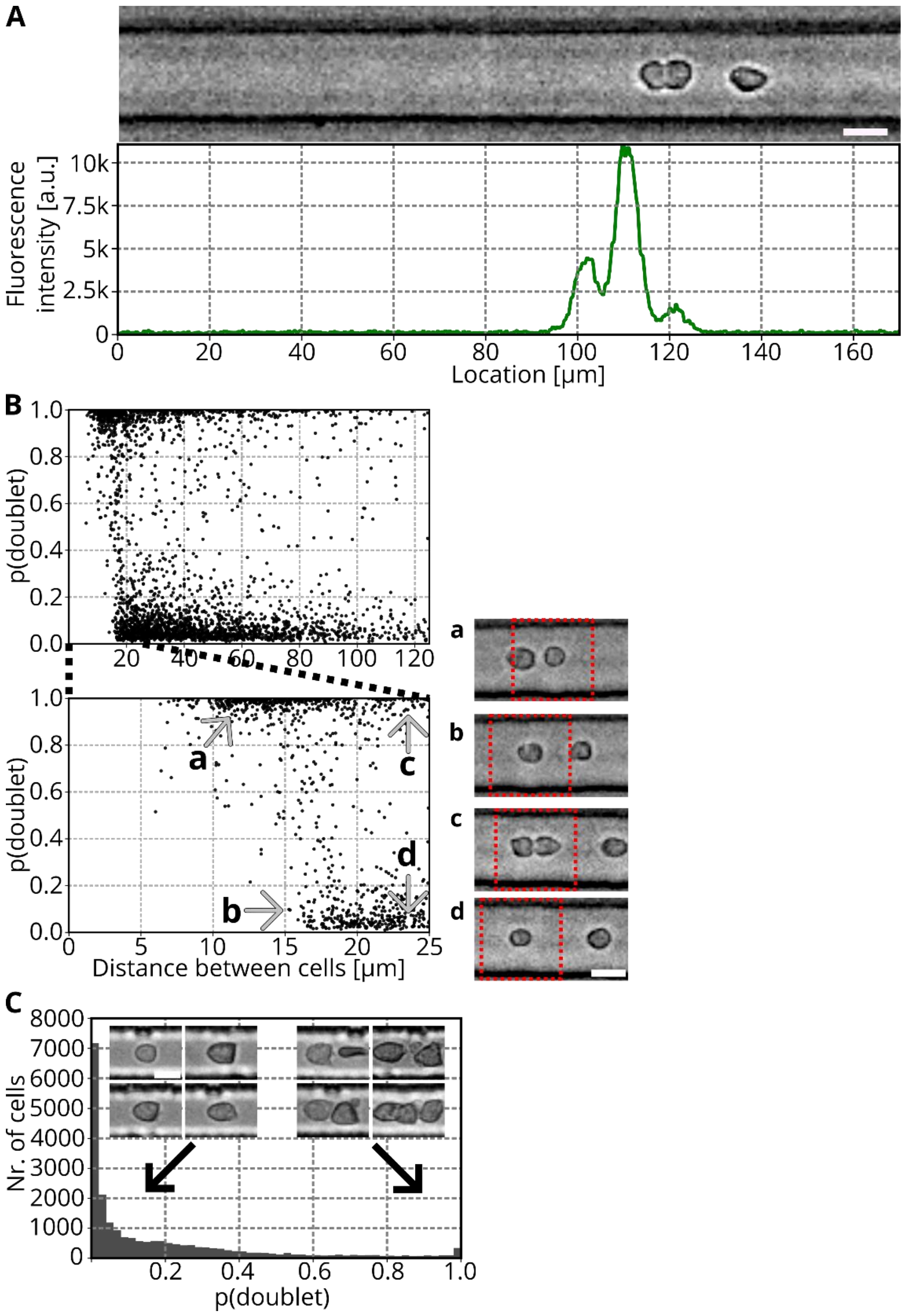


Figure S3: Detection of cell aggregates

(A) Example event of an RT-FDC measurement of Nrl-eGFP. Image on the top shows the captured bright-field image, with one doublet, closely followed by another cell. The plot below shows the corresponding fluorescence track, indicating that assigning fluorescence values to each cell would be difficult. Scale bar: 10 µm.

(B) Leading/ trailing cell detection. In the dataset, events were labeled as doublet if a second cells was closer than 15 µm. The scatterplot shows the probability to be a doublet p(doublet) vs. the absolute distance between cells. Especially, the zoomed-in version (below) shows that the 15 µm threshold was adopted by the model. Four regions (a-d) are marked in the plot and example images are shown. The model input image size is 36x36 pixels (=24.5x24.5 µm) and this region is indicated in each image by a red rectangle. Scale bar: 10 µm

(C) RT-FDC measurement of blood cells. Histogram shows the probability of events to be a doublet (p(doublet)). Example images show the phenotype for low and high p(doublet). The figure shows the same measurement as Figure 2C. Scale bar: 10 µm.

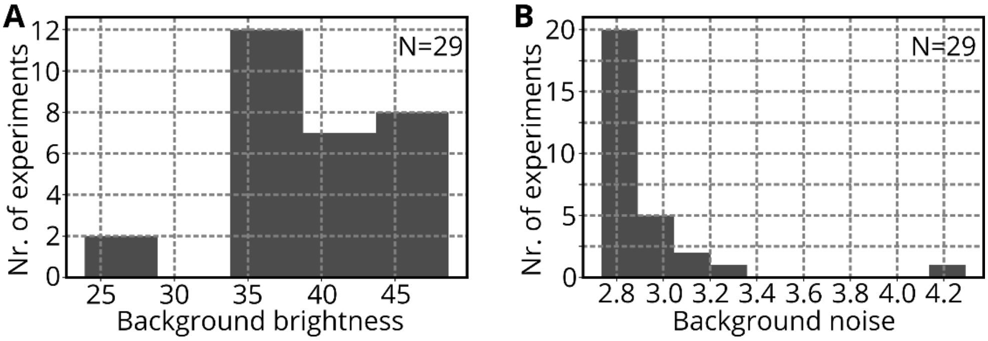


**Figure S4: Range of brightness and noise levels in images**

**(A) Histogram shows the median background brightness values of 29 experiments. Background brightness values were obtained by computing the median of the grayscale levels of the pixels in a region (10x255 pixels), located above the measurement channel.**

**(B) Histogram shows the standard deviation values of 29 experiments. Standard deviation values were obtained by computing the standard deviation of the grayscale level of the pixels in a region (10x255 pixels), located above the measurement channel.**

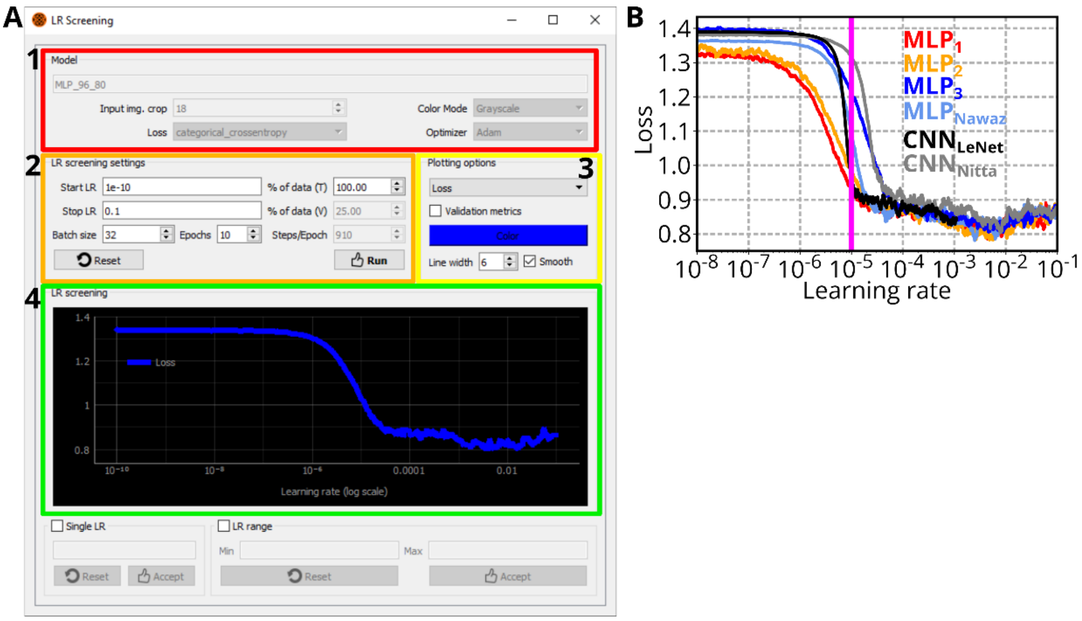


Figure S5: Learning rate screening

(A) Screenshot of the learning rate screening window in AIDeveloper. Colored rectangles in the image indicate the workflow:

1 – Information of the selected model

2 – Settings for the learning rate screening. Start and Stop LR indicate the range for the screening. Optionally, the screening can be performed using only a certain amount of the training data ("% of data (T)"). During a screening, the model is trained for some iterations (called "Epochs"). The button "Run" initiates the screening.

3 – During the screening, loss, accuracy and the first derivative of both are tracked. Using the dropdown menu, each of those can be plotted. Furthermore, options to customize the plot are available.

4 – Plotting region. After a right-click the user has the option to export the plot as image or the underlying data as hd55 or excel file.

(B) Result of the learning rate screening for all MLPs and CNNs. During screening, models are trained using the available training data and data augmentation methods are applied. The learning rate screening was performed using AIDeveloper 0.2.3.

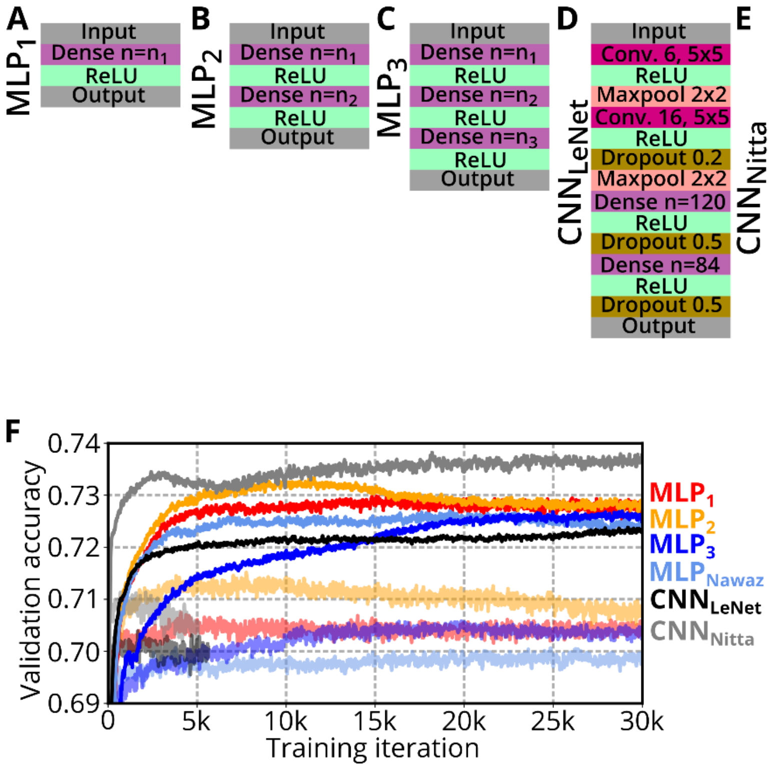


Figure S6: Model architectures and training
(A)-(E) Sketches show the model architectures of multilayer perceptron with one, two and three hidden layer and convolutional neural net with two and four convolutional layers.
(F) Plot shows the validation accuracy of models trained for 30,000 iterations. Models were trained using the learning rate $10^{-5}$ (opaque lines) or $10^{-3}$ (transparent lines).

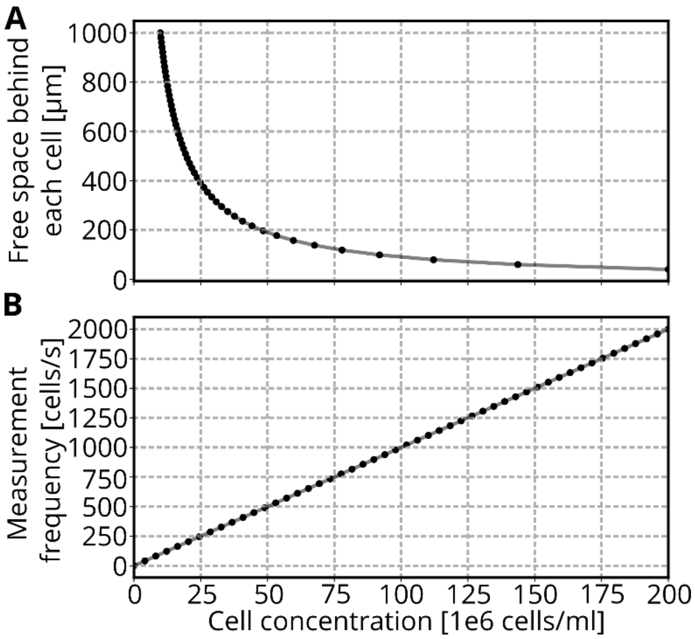


**Figure S7: Cell concentration vs. space between cells and measurement frequency**

**(A) Scatterplot shows the calculated free space behind each cell in the measurement channel at various cell concentrations (in the sample syringe). The calculation assumes a homogenous concentration profile and a diameter of the cell of 10 µm.**

**(B) Scatterplot shows the calculated measurement frequency, based on the assumption that cells are homogenously distributed in the sample volume and move at a flow rate Q=0.04 µl/s through the chip (sample flow rate: $Q_{sample}$=0.01 µl/s, sheath flow rate: $Q_{sheath}$=0.03 µl/s):** $Measurement\ freq. = Q \cdot cell\ concentration$

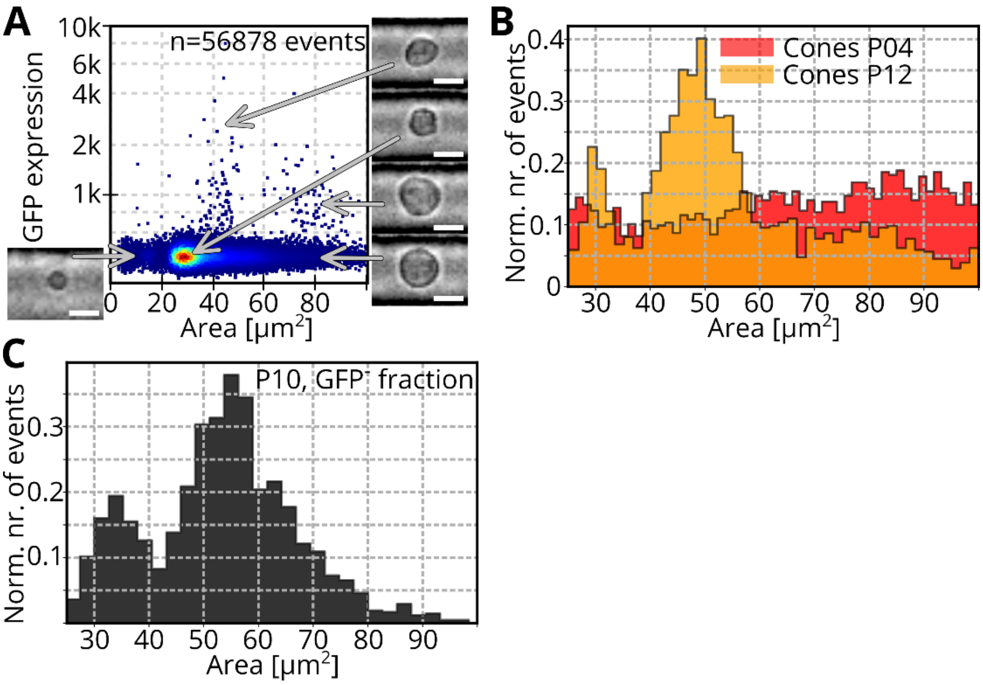


Figure S8: Other cell types
(A) Scatterplot shows an RT-FDC measurement of mGluR6-GFP bipolar reporter mouse retina cells at P04. The $GFP^+$ bipolar cells are rare (0.38%) which can be explained since the marker is only expressed in a subset. $GFP^+$ bipolar cells are mostly located in a size region around 40 $\mu m^2$. Inset images show example cell phenotypes I certain regions of the plot. Scale bars: 10 μm.
(B) RT-FDC measurement of cone-GFP mouse retina cells at P04 and P12. The histogram shows the area distributions of the $GFP^+$ fraction. For both cases, approximately 1.4% of the events were $GFP^+$.
(C) Histogram shows the $GFP^-$ fraction (FAC-sorted) of an Nrl-eGFP mouse at P10. At this maturation stage, no retinal progenitor cells are remaining. However, there are still cells in the same size region like rod photoreceptors. Data taken from [1].

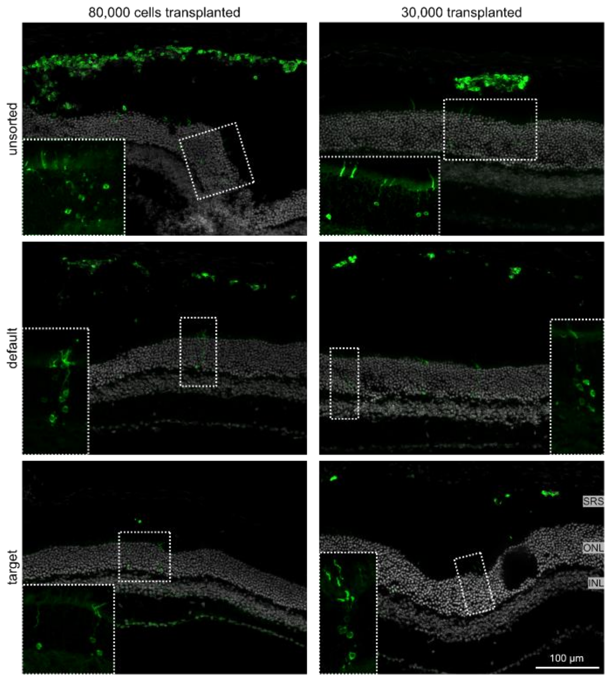


Figure S9: Transplantation results from unsorted, default, and target fractions
Exemplary images of recipient mouse retina collected 2 weeks after transplantation of 80,000 or 30,000 cells per eye. In addition to the target fraction, unsorted cells, and cells collected in the soRT-DC default outlet were used as controls. With all conditions allowing detection of Nrl-eGFP$^+$ cells in the subretinal space and eGFP$^+$ cytosol in the recipient retina (magnifications), the “unsorted” eyes clearly contain more eGFP$^+$ cells than both the “default” and the “target” eyes. SRS = subretinal space; ONL = outer nuclear layer; INL = inner nuclear layer.
